# Defining a tandem repeat catalog and variation clusters for genome-wide analyses

**DOI:** 10.1101/2024.10.04.615514

**Authors:** Ben Weisburd, Egor Dolzhenko, Mark F. Bennett, Matt C. Danzi, Isaac R. L. Xu, Hope Tanudisastro, Bida Gu, Adam English, Laurel Hiatt, Tom Mokveld, Guilherme De Sena Brandine, Readman Chiu, Nehir Edibe Kurtas, Helyaneh Ziaei Jam, Harrison Brand, Indhu Shree Rajan Babu, Melanie Bahlo, Mark JP Chaisson, Stephan Züchner, Melissa Gymrek, Harriet Dashnow, Michael A. Eberle, Heidi L. Rehm

## Abstract

Tandem repeat (TR) catalogs are important components of repeat genotyping studies as they define the genomic coordinates and expected motifs of all TR loci being analyzed. In recent years, genome-wide studies have used catalogs ranging in size from fewer than 200,000 to over 7 million loci. Where these catalogs overlapped, they often disagreed on locus boundaries, hindering the comparison and reuse of results across studies. Now, with multiple groups developing public databases of TR variation in large population cohorts, there is a risk that, without sufficient consensus in the choice of locus definitions, the use of divergent repeat catalogs will lead to confusion, fragmentation, and incompatibility across resources.

In this paper, we compare existing TR catalogs and discuss desirable features of a comprehensive genome-wide catalog. We then present a new, richly annotated catalog designed for large-scale analyses and population databases. This new catalog, which we call the TRExplorer catalog v1.0, contains 4.86 million TR loci and, unlike most catalogs, is designed to be useful for both short-read and long-read analyses. It consists of 4,803,366 STRs and 59,675 VNTRs, of which 780,607 STRs and 21,888 VNTRs are both polymorphic and entirely absent from widely-used catalogs previously developed for short-read analyses. Additionally, our catalog stratifies TRs into two groups: 1) isolated TRs suitable for repeat copy number analysis using short-read or long-read data and 2) so-called variation clusters that contain TRs within wider polymorphic regions that are best studied through sequence-level analysis. To define variation clusters, we present a novel algorithm that leverages long-read HiFi sequencing data to group repeats with surrounding polymorphisms. We show that the human genome contains at least 25,000 complex variation clusters, most of which span over 120 bp and contain five or more TRs. Resolving the sequence of entire variation clusters instead of individually genotyping constituent TRs leads to a more accurate analysis of these regions and enables us to profile variation that would have been missed otherwise. We also share the <u>trexplorer.broadinstitute.org</u> portal which allows anyone to search, visualize, and download the catalog along with variation clusters and annotations.

## Introduction

Tandem repeats (TRs) are nucleotide sequences composed of consecutive repetitions of a shorter motif (also called a repeat unit). Typically, TRs with motif size of 6 bp or less are called short tandem repeats (STRs) while those with longer motifs are called variable number tandem repeats (VNTRs). These sequences can be perfect repeats of a single motif–for example, CAG CAG CAG CAG–or they can contain interruptions, such as CAG CAG CA**A** CAG. Over the past fifteen years, wide-ranging studies have used sequencing data to investigate many aspects of TRs including mutation rates,^1–4^ effects on gene expression,^5^ splicing,^6^ methylation,^7,8^ and disease risk.^9–11^ With the exception of several specialized methods for identifying *de novo* variation,^12–14^ most TR genotyping tools^15–24^ involved in these studies require the user to provide a TR catalog which specifies the reference coordinates and motifs of each TR locus to genotype–making the choice of catalog a key factor in the overall sensitivity of a study.^25–27^

Researchers have typically generated TR catalogs using tools like Tandem Repeats Finder (TRF)^28^ that detect repetitive sequences by scanning the reference genome. Although a reference-based approach is able to identify a wide range of TR loci, it can miss polymorphic repeats that are too short or absent from the reference genome. Furthermore, methods like TRF are highly sensitive to input parameters, with some commonly-used settings yielding approximately 1 million, mostly perfect TRs,^16^ while other settings produce over 10 million TRs (**Table S1**) that contain many sequence interruptions.^29^ An alternative approach, which we call the cohort-driven strategy, detects TR variation directly from sequencing data^12,13,30^ or from haplotype-resolved assemblies.^29,31^ The cohort-driven strategy not only prioritizes polymorphic loci, but can also capture TRs that are not well represented in the reference genome and thus missed by the reference-based approach.

In addition to the different strategies for identifying TR loci, another important aspect of catalog design is the definition of locus boundaries. Since naturally occurring repeat sequences can contain interruptions or consist of multiple arrays of highly similar motifs, defining the starts and ends of TR loci in the reference genome is often subject to ambiguity. Yet, even small, seemingly inconsequential differences in how locus boundaries are specified within different catalogs, resources, or reference materials can lead to real issues with interpretation of TR genotypes. For example, the *PABPN1* locus has two commonly used definitions. With narrower locus boundaries counting only perfect GCG motifs, the pathogenic threshold is 8 GCG repeats, while with wider boundaries that allow GCA interruptions, the equivalent pathogenic threshold is 12 repeats.^32^ This creates the potential for inadvertent errors in the diagnosis of autosomal dominant oculopharyngeal muscular dystrophy (OPMD) if the genotype (number of repeats) in a given individual is interpreted without awareness of which locus definition was used during genotyping (**Supplementary Information**).

The selection of TR loci and the specification of their boundaries can influence the suitability of a catalog for two different kinds of downstream analyses. The first approach focuses primarily on the quantification of repeat copy numbers and using them to detect outlier expansions or perform association tests with other variables like gene expression. The second approach involves sequence-level analysis of TR regions in order to characterize motif composition, interruption patterns, and properties of flanking sequences. Most existing TR genotyping tools, especially those designed for short-read data, only report repeat copy numbers and not allele sequences.^15,16^ They are unable to resolve the full sequence-level variations of complex regions such as nested repeats and repeats surrounded by other repeats or structural variants. Additionally, these tools work best with narrow locus definitions that contain only perfect or nearly-perfect repeat sequences.^31^ Therefore, catalogs designed for repeat copy number analysis benefit from setting narrow locus boundaries, and flagging TRs within complex regions as more likely to produce erroneous genotypes. On the other hand, the latest generation of tools^17,18,23,24^ can report allele sequences in addition to repeat copy numbers, and so enables sequence-level analysis both for isolated TRs as well as more complex repeat regions.^33,34^ Examples of TR variation that especially benefit from sequence-level analysis include the recently identified polymorphic region associated with the stability of the *FGF14* repeat,^35^ a single base pair insertion in a homopolymer located within a *MUC1* VNTR that causes autosomal dominant tubulointerstitial kidney disease (ADTKD),^36^ and TRs like the *RFC1* locus whose pathogenicity is affected by sequence composition changes.^37,38^ Such loci frequently correspond to short stretches of perfect tandem repeats punctuated by insertions, deletions, and substitutions in and around the repeat sequences.

Taken together, these observations highlight the challenges of creating genome-wide TR catalogs suitable for population studies, especially multi-center studies employing different computational tools and sequencing technologies. To help address these challenges, we introduce a new genome-wide repeat catalog. We employ both cohort-based and reference-based TR identification approaches to capture a comprehensive set of loci, including those that harbor common or rare variation. To simplify adoption, we share the catalog in the different formats used by popular TR analysis tools for short- and long-read data. Our catalog also stratifies TRs into two groups: isolated TRs where repeats are surrounded by non-polymorphic flanking regions and variation clusters that contain TRs surrounded by polymorphic sequence. Due to their complexity, variation clusters are best studied through sequence level analysis, particularly using long-read data. To perform this stratification we introduce a new method for profiling population-scale variation around TRs and other regions of the genome.

Lastly, we provide the trexplorer.broadinstitute.org portal to enable interactive online exploration of the TRs and variation clusters in our catalog. Users can view population allele frequency distributions, visualize locus definitions from different catalogs, and filter TRs by various criteria such as gene region, repeat motif, and/or polymorphism rates. Users can also export the full catalog or a filtered subset to different file formats suitable for downstream analysis.

## Results

### Existing tandem repeat catalogs

Widely-used TR catalogs differ substantially in their core attributes such as their total number of loci, range of included motif sizes, repeat purity, and distribution of repeat sizes in the reference (**Table 1**, **Figure 1**, **Figure S1, Figure S2, Methods**). Strikingly, pairwise comparison of eleven TR catalogs showed that every pair differed in over half of their TR definitions (**Figure S1**). These catalogs were generated using different analysis strategies (reference-based or cohort-driven) and aimed at different study objectives, tools and sequencing technologies. This led to different minimum repeat length and purity thresholds choices when selecting TR loci for inclusion (**Table S1**). Also, the repeat cataloging projects employed distinct approaches to define locus boundaries. For example, the recently-released Adotto v1.2^39^, Platinum TRs v1.0^4^, and the Chiu et al. 2024^29^ catalogs contain many long imperfect repeats and include non-repetitive flanking sequences. Such locus definitions are compatible with existing tools for genotyping TRs in long reads, but often cannot be accurately genotyped with existing tools for short-read sequencing data. On the other hand, older catalogs like GangSTR v17 (1.3 million loci) and Illumina 174k (174 thousand loci) define narrow locus boundaries around mostly perfect TR sequences and so are suitable for both short-read and long-read analyses. However, for technical or historical reasons, these early catalogs are smaller than more recent catalogs and, importantly, are missing more than half of our test set of 250,262 orthogonally-derived polymorphic STR loci with 3-6 bp motifs (**Methods**).

**Figure 1.**
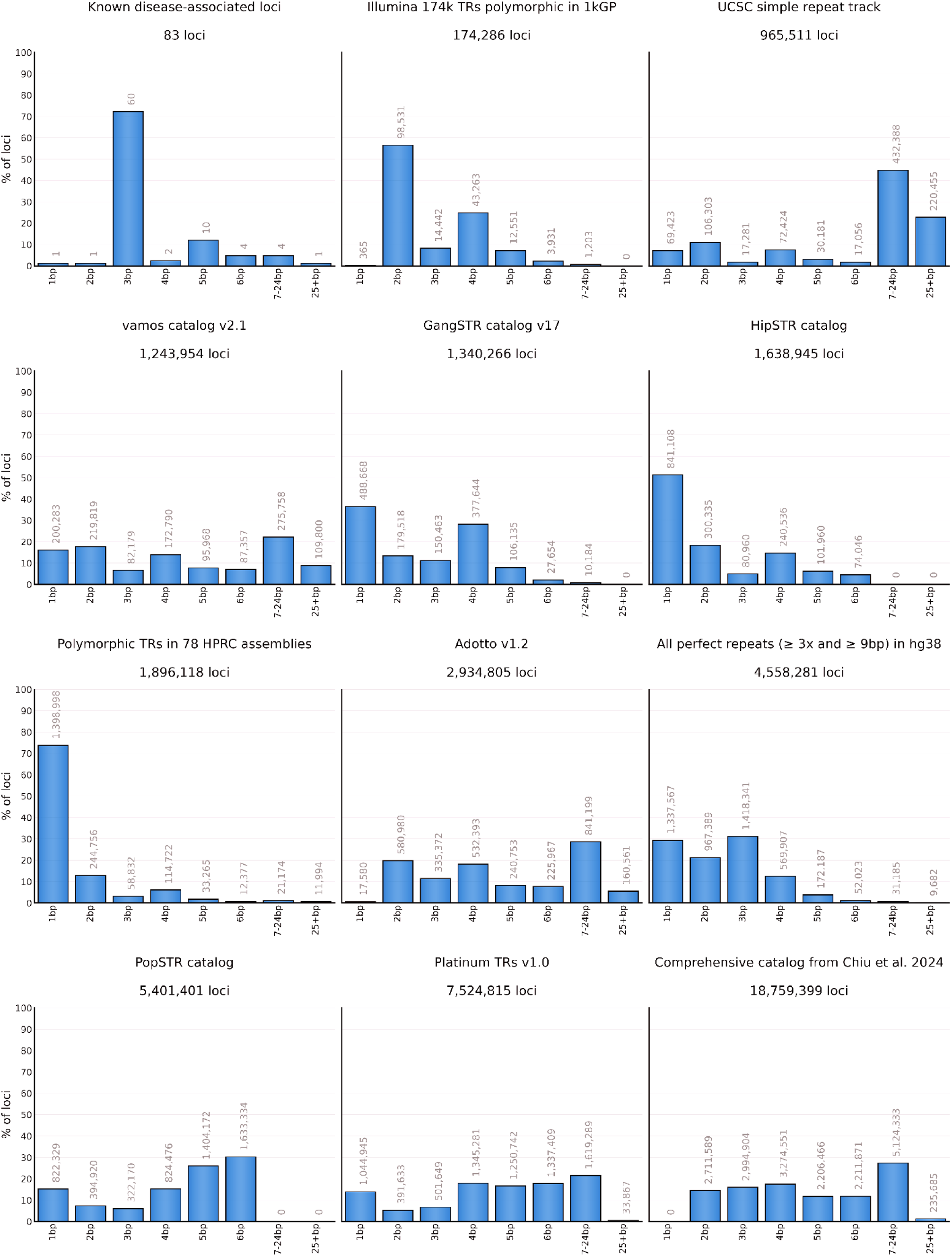
Comparison of motif size distributions across TR catalogs. The x-axis represents motif sizes. Each bar is labeled with the number of loci it contains.

**Table 1:** Properties of commonly-used TR catalogs, including the total size, total span, and motif size ranges. The “Polymorphic STRs missed” column is based on lack of overlap with a test set of 250,262 polymorphic STRs with 3-6 bp motifs identified in 78 diverse haplotype-resolved assemblies. Overlap was defined leniently as having any TR definition that overlapped the test set STR by 1 bp or more, regardless of motif. * For the Adotto v1.2 catalog, each of the original 1,784,804 entries was split into its constituent TRs, before calculating statistics ** The PlatinumTR v1.0 catalog was converted from TRGT catalog format, splitting any compound locus definitions into their constituent TRs (**Methods**). † 296,782 loci (3.8%) were excluded because their reference sequences contained interruptions that prevented us from unambiguously splitting the region into its constituent repeats.

| row | catalog | total loci | genome coverage | motif sizes | locus sizes in reference | pure repeat fraction in reference | includes chrX | includes chrY | polymorphic STRs missed |
| --- | --- | --- | --- | --- | --- | --- | --- | --- | --- |
| 1 | Known disease-associated TRs + adjacent repeats & historical candidate loci | 83 | <0.01% | 1 - 27 bp | 6 - 150 bp | 94% | ✓ | ✗ | N/A |
| 2 | Illumina 174k polymorphic loci | 174,286 | 0.15% | 1 - 24 bp | 4 - 532 bp | 100% | ✓ | ✗ | 71% |
| 3 | UCSC Simple Repeat Track | 965,511 | 9.9% | 1 - 1,991 bp | 13 - 499,998 bp | 83% | ✓ | ✓ | 38% |
| 4 | Vamos v2.1 | 1,243,954 | 2.9% | 1 - 667 bp | 1 - 9,975 bp | 64% | ✓ | ✓ | 27% |
| 5 | GangSTR v17 | 1,340,266 | 0.74% | 1 - 20 bp | 10 - 612 bp | 100% | ✓ | ✓ | 53% |
| 6 | HipSTR | 1,638,945 | 1.4% | 1 - 6 bp | 9 - 132,210 bp | 77% | ✓ | ✓ | 30% |
| 7 | Adotto v1.2 * | 2,934,805 | 3.9% | 1 - 495 bp | 4 - 49,802 bp | 85% | ✓ | ✓ | 13% |
| 8 | popSTR | 5,401,401 | 2.7% | 1 - 6 bp | 8 - 136 bp | 96% | ✗ | ✗ | 26% |
| 9 | Platinum TRs v1.0 ** | † 7,524,815 | 5.1% | 1 - 822 bp | 4 - 9,504 bp | 90% | ✓ | ✓ | 45% |
| 10 | Chiu et al. 2024 | 18,759,399 | 10.7% | 2 - 100 bp | 2 - 185,860 bp | 70% | ✓ | ✗ | 4% |

### Defining a genome-wide catalog

To incorporate the advantages of both reference-based and cohort-driven approaches, we collected loci from three cohort-driven catalogs and one reference-based catalog (**Table 2**). Where multiple source catalogs contained different definitions of the same TR locus (**Methods**), the higher prioritized definition–as described in Table 2–was incorporated. First, we included 63 disease-associated loci, 10 of their adjacent repeats, as well as 10 candidate TR loci that have been historically included in rare disease analyses. These loci have been the focus of extensive research,^40,41^ and many have well established, widely-used locus definitions which we incorporated here to ensure compatibility with prior studies. Next, we added 174,244 loci from the Illumina catalog of polymorphic TRs which was created in 2021 by analyzing signals of TR variation within short-read sequencing data from 2,504 individuals in the 1000 Genomes Project.^30^ Prioritizing this catalog above catalogs 3 and 4 (**Table 2**) ensured compatibility with the locus boundary definitions used in prior studies and resources.^42^

**Table 2:** Four source catalogs that were incorporated into our tandem repeat catalog.

| priority | catalog | total loci | genome coverage | motif sizes | locus sizes in reference | pure repeat fraction in reference | includes chrX | includes chrY | loci contributed to final catalog |
| --- | --- | --- | --- | --- | --- | --- | --- | --- | --- |
| 1 | Known disease-associated TRs + adjacent repeats & historical candidate loci | 83 | <0.01% | 1 - 27 bp | 6 - 150 bp | 94% | ✓ | ✗ | 83 |
| 2 | Illumina 174k polymorphic TRs in 1kGP | 174,286 | 0.15% | 1 - 24 bp | 4 - 532 bp | 100% | ✓ | ✗ | 174,244 |
| 3 | Perfect repeats that span $\geq 9$ bp and $\geq 3$ repeats in hg38 | 4,558,281 | 2.0% | 1 - 833 bp | 9 - 2,523 bp | 100% | ✓ | ✓ | 4,391,197 |
| 4 | Polymorphic TRs in 78 T2T assemblies | 1,937,805 | 1.0% | 1 - 833 bp | 1 - 2,499 bp | 100% | ✓ | ✓ | 297,517 |
|  | Tandem Repeat Catalog | 4,863,041 | 2.1% | 1 - 833 bp | 1 - 2,523 bp | 100% | ✓ | ✓ |  |

To ensure comprehensive inclusion of perfect repeat sequences, we identified all TRs in hg38 that spanned ≥3 copies of any 3-1000 bp motif, ≥5 copies of a dinucleotide motif, or ≥9 copies of a homopolymer motif (**Methods**). This added another 4,391,197 loci to the catalog. Our focus on perfect repeats was motivated by their higher mutation rates compared to interrupted repeats, the importance of repeat purity in TR-associated disease mechanisms,^43,44^ as well as their higher overall genotyping accuracy in short-read analyses.^16^ We chose a ≥ 3x threshold to match several known disease-associated loci such as *C9orf72* that span only three perfect repeats in the hg38 reference. Additionally, our initial analyses showed that this threshold represented a favorable balance between precision and recall, capturing many polymorphic TR loci in our test set without excessively inflating the size of the catalog (Figure S2 in Weisburd et al. 2023).^31^

Subsequently, to identify the longest pure segments (LPS) we applied TRGT followed by TRGT-LPS (**Methods**) to long-read samples from 1,027 individuals in the AoU phase 1 release as well as 256 individuals from the Human Pangenome Reference Consortium (HPRC) in order to directly measure the polymorphism rates at 4,023,736 out of 4,391,197 of these loci. This analysis showed that only 753 out of 4,023,736 loci (0.019%) were non-polymorphic (i.e. homozygous reference) in all individuals. However, because genotyping errors can artificially inflate the number of polymorphic loci, we also evaluated the standard deviation of the allele size distribution at each locus and found that the standard deviation exceeded 0.1 in the AoU phase 1 or the HPRC samples at 2,924,712 out of 4,023,736 (73%) loci, while at 1,446,103 (36%) loci it exceeded 0.3 which is higher than the standard deviation of some known disease-associated STR loci such as ZIC3 and PABPN1 (**Table S2**). We kept low polymorphism repeats in our catalog to enable studies aimed at profiling variation in populations that were not well represented in our data, to allow different studies to set their own minimum polymorphism thresholds when selecting loci for inclusion, and finally to differentiate these loci from ones that were absent from our polymorphism analyses.

Finally, we incorporated polymorphic TRs detected using 78 haplotype-resolved T2T assemblies from the Human Pangenome Reference Consortium (HPRC) and Human Genome Structural Variation Consortium (HGSVC). The computational method we employed for this task (**Methods**) allowed us to complement the previous three catalogs by capturing polymorphic loci that had fewer than three repeats in the reference, or that contained single base interruptions like those observed at known disease-associated loci such as *ZIC3*, *ATXN1, and FXN*. This approach contributed a further 297,519 TRs to the final catalog. It is worth noting that our catalog covers all loci in the GangSTR catalog (v17) even though we did not explicitly include it as a source catalog.

The resulting TRExplorer v1.0 catalog contained 4,863,041 TRs that collectively spanned 2.1% of the hg38 reference. As summarized in **Figure S3**, 4,803,366 of these (98.8%) were STRs, including 1,567,337 homopolymers (32.2%), 978,972 dinucleotide repeats (20.1%), 1,432,117 trinucleotide loci (29.4%), 590,787 loci with 4bp motifs (12.1%), 177,422 loci with 5bp motifs (3.6%), and 56,731 loci with 6bp motifs (1.2%). 59,675 were VNTRs (1.2%). Intersecting the catalog with Gencode v49 gene annotations and taking the most significant gene region based on the following priority: coding region (CDS), 5’ untranslated region (UTR), 3’ UTR, exon, intron, promoter, showed that 3,200,611 (65.8%) TRs in the catalog were intronic, 1,353,371 (27.8%) were intergenic, 49,496 (1.0%) overlapped promoters, 64,929 overlapped 3’ UTRs (1.3%), 125,242 (2.6%) overlapped exons of non-coding genes, 41,420 (0.9%) overlapped coding regions, and 27,972 (0.6%) overlapped 5’ UTRs.

To estimate the polymorphism rates of TR loci in the catalog, we used TRGT followed by TRGT-LPS (**Methods**) to genotype 256 HPRC HiFi samples from diverse ancestries, as well as the AoU phase 1 release data from 1,027 HiFi samples. In aggregate, 2,438,741 (52%) loci were polymorphic, having an allele size distribution standard deviation ≥ 0.2 in one or both of these datasets (**Figure S4A**). Stratifying polymorphism rates by motif indicated trinucleotide repeats have the highest fraction of non-polymorphic loci (**Figure S4B**). Stratifying by source catalog (Table 2, **Figure S4C**) showed that the Illumina 174k catalog had the highest fraction of polymorphic loci (98%), followed by the catalog of known disease-associated loci (84.3%), followed by the catalog of polymorphic TRs in T2T assemblies (72.6%), followed by perfect repeats in hg38 that span ≥ 9bp and ≥ 3 repeats (48.4%). Downsampling analysis in individuals of African ancestry from the HPRC data showed that, after the first 50 samples, every additional sample increased the total number of loci with more than one observed allele size by 4.5k on average, while the number of loci with more than 2 observed allele sizes increased by 3.3k loci, and there was no indication that the number of polymorphic loci plateaued after 90 samples (**Figure S4D**).

### Analysis of variation clusters

Some tandem repeats (TRs) are located within regions that have high rates of polymorphism and hence often differ significantly from the reference genome (**Figure 2 A-C**). We refer to such regions as variation clusters (VCs). Because most methods for TR analysis assume that the regions surrounding the TR closely resemble the reference genome, TRs located within variation clusters may be prone to analysis errors. For example, consider (AT)n repeats located in a variation cluster in the *KCNMB2* gene (**Figure 2D**). In the individual shown, the flanks of these TRs significantly differ from the reference genome, making it difficult to locate the exact boundaries or separate one repeat from the other repeats located within this region. This issue can be avoided if we locate the boundaries of the entire VC and then determine its full-length allele sequences in each sample with tools like TRGT^18^ or LongTR.^17^ Then, the resulting sequences can be either analyzed directly or their repeat content can be characterized with existing methods,^18^ making it possible to analyze VCs together with isolated TRs.

**Figure 2:**
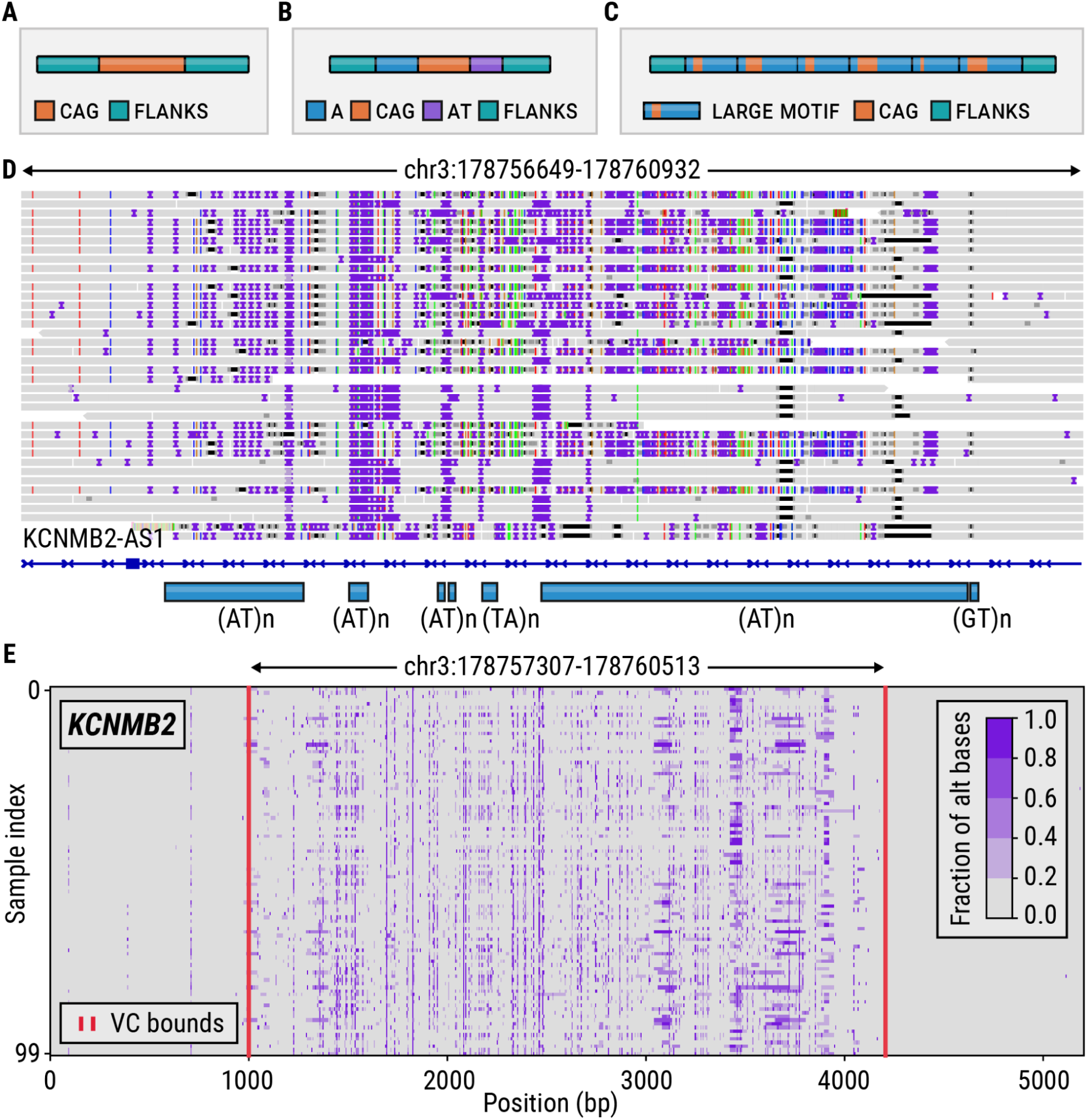
(A) A TR surrounded by non-repetitive flanks. (B) A TR with a CAG motif surrounded by an A homopolymer and AT dinucleotide repeat. (C) Nested TRs. (D) HiFi reads from the HG002 sample spanning a variation cluster located in an intron of the *KCNMB2* gene. (E) A variation plot depicting the *KCNMB2* locus in 100 long-read HiFi samples; the red vertical lines denote the boundaries of variation cluster (VC) computed by the vclust tool.

We developed a method, vclust, that when given a set of TR regions and aligned sequencing reads from multiple samples can identify variation clusters around each TR (**Methods**). We first used vclust to characterize variation across 63 loci containing known disease-associated TRs in 100 HiFi WGS HPRC samples. Our method identified variation clusters around eight of these TRs located in the *ATXN8OS*, *FGF14*, *NOP56*, *CNBP*, *HTT*, *BEAN1*, *LRP12*, and *EIF4A3* genes. Variation around most of these TRs has already been characterized: the *ATXN8OS*, *NOP56*, *CNBP*, and *HTT* TRs contain known adjacent repeats that contribute to variation in their flanks. A polymorphic region flanking the *FGF14* TR was recently found to have a stabilizing effect on this repeat.^35^ As anticipated, the vclust analysis incorporated this region into the *FGF14* variation cluster (**Figure 3A**). The *CNBP* VC (**Figure 3B**) includes three polymorphic repeats: CAGG, CAGA, and CA motifs while the *NOP56* VC (**Figure 3C**) contains the original repeat, plus a common deletion located less than 50 bp away from the repeat boundary. *BEAN1*, *LRP12*, and *EIF4A3* VCs were caused by the presence of small indels adjacent to these repeats.

**Figure 3.**
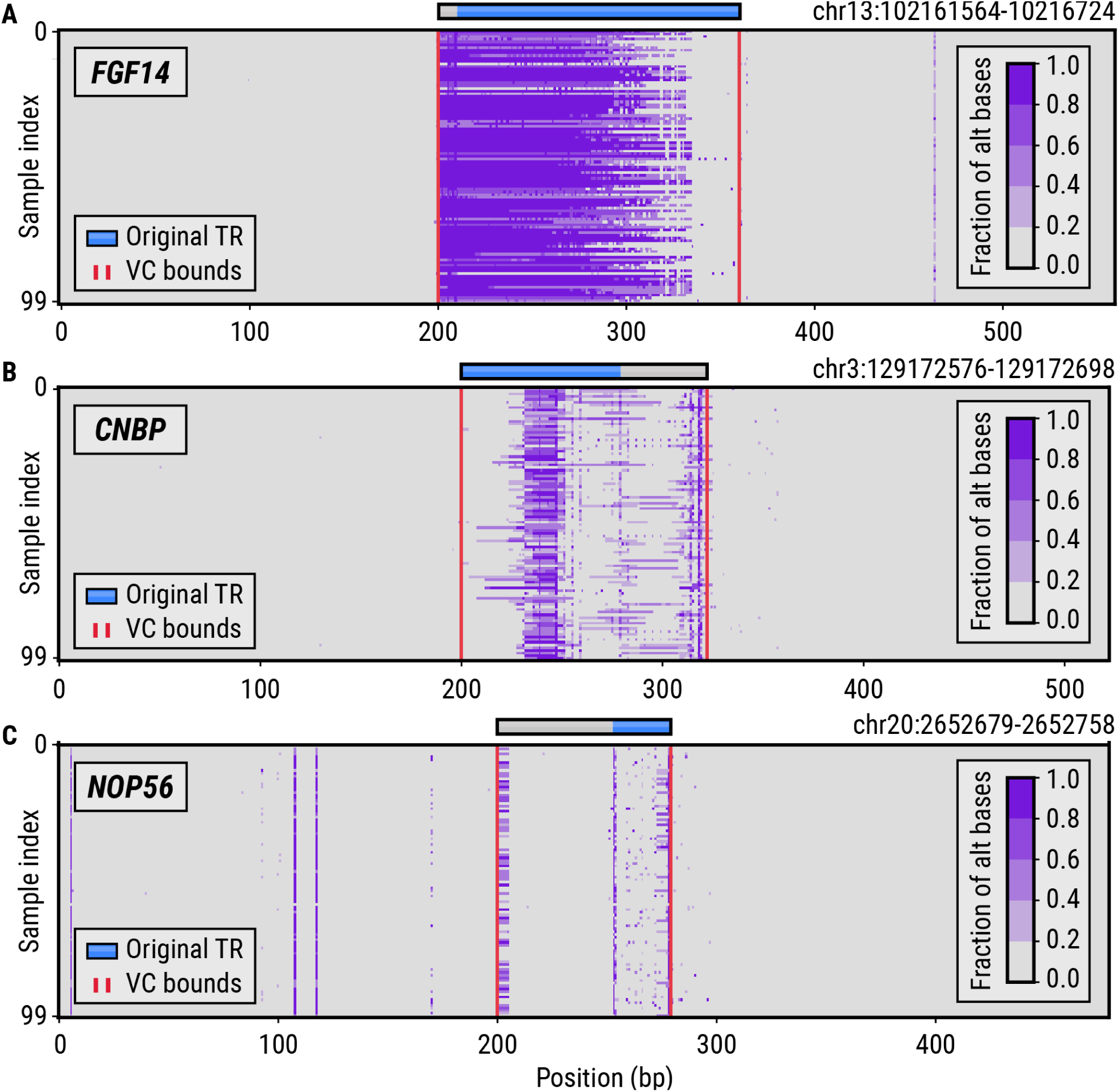
Variation plots showing the genetic variation in 100 HiFi HPRC samples around known pathogenic repeats in the (A) *FGF14*, (B) *CNBP*, and (C) *NOP56* genes. The blue horizontal bars denote the original repeat region while the gray bars depict the extension of the repeat region to a full-length variation cluster. The red vertical lines denote the boundaries of variation clusters (VCs) computed by the vclust tool.

Next, we applied the variation cluster analysis to our catalog of 4,863,041 TRs. This analysis identified 273,112 VCs that collectively overlapped 744,458 TRs, with 66% of VCs overlapping two or more TRs. About 18% of VCs (N=50,557) spanned more than 120 bp in the reference genome (**Figure 4A**), a length at which accurate genotyping with short-read data (30x 150bp WGS) becomes difficult. To assess the benefits of genotyping entire variation clusters rather than their constituent TRs, we compared the accuracy of allele sequences obtained by genotyping full VCs with those obtained by genotyping individual TRs in a long-read HiFi sequencing dataset from the HG002 sample. Accuracy was defined as sequence-level concordance with a highly accurate assembly of the same genome^45,46^ (**Methods; Assembly consistency analysis**). The allele concordance rate increased from 79.43% for TRs (87.30% when off-by-one repeat errors were considered concordant) to 90.58% (95.98%) for the VCs containing these TRs. In contrast, isolated TRs showed a concordance rate of 96.21% (98.50%), indicating that TRs located within VCs are a major source of genotyping errors when analyzed individually. To further characterize VCs containing multiple repeat tracts, we identified 24,867 VCs that each contained five or more constituent TRs, which we refer to as complex VCs. These complex VCs collectively contain 254,879 TRs, have a median reference length of 208 bp, and in 86% of cases exceed 120 bp (**Figure 4B**). For example, a VC in the *KCNMB2* gene (**Figure 2E**) spans over 3 Kbp in the reference genome and contains 135 TRs, including 116 TRs with AT motifs. Other similarly complex regions can benefit from VC analysis (**Figure S5**).

**Figure 4.**
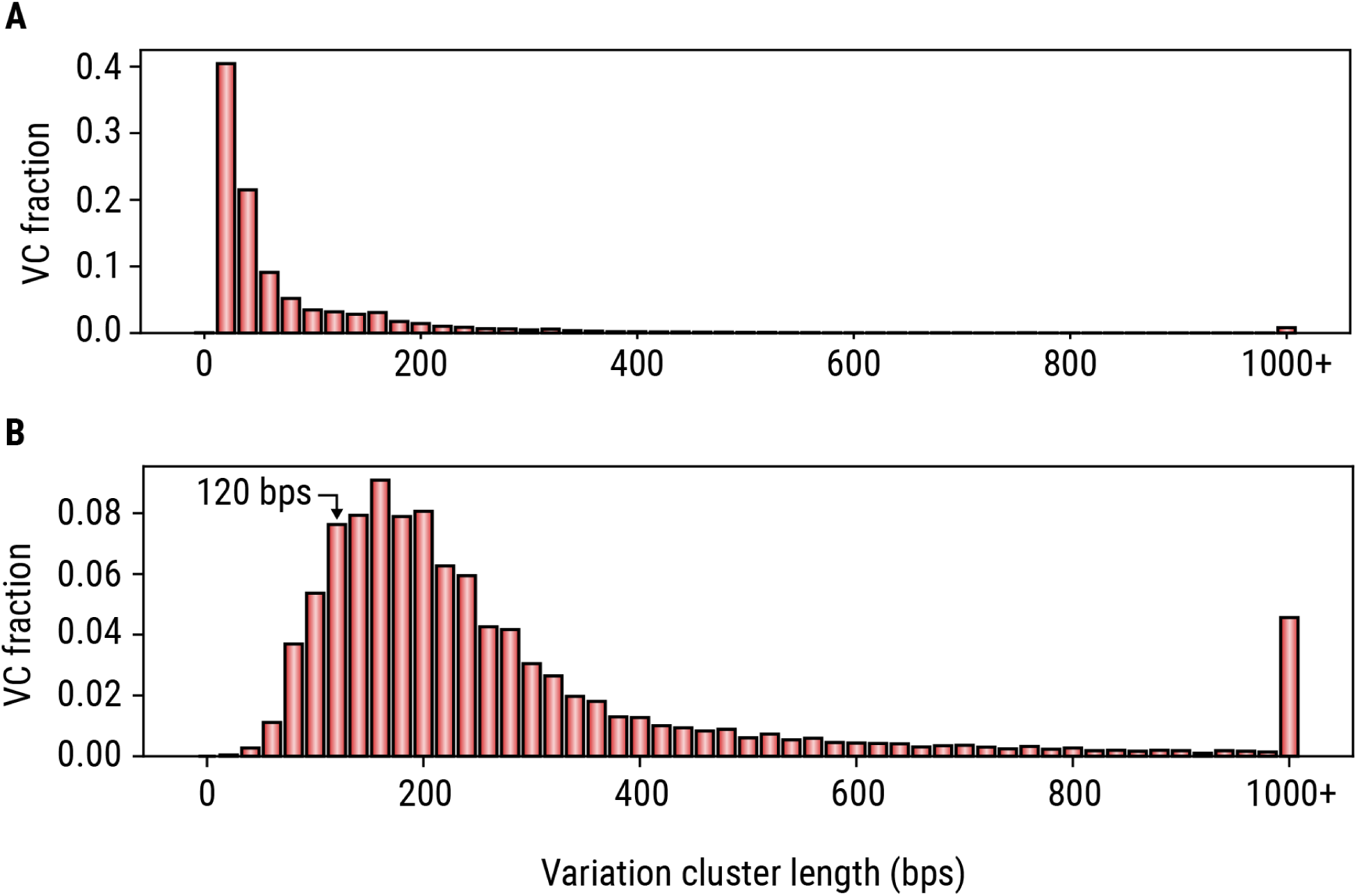
Size distributions of variation clusters. (A) Distribution of the differences between variation cluster length and the length of the original TR region contained within it. (B) Distribution of lengths of complex variation clusters in the reference genome.

We investigated the distribution of features that differed between isolated TRs compared to those within VCs or complex VCs (**Figure 5**). The majority of TRs with 1-9 bp repeat motifs are isolated TRs (88%), with this trend being most pronounced for homopolymer and trinucleotide TRs (93% and 94%, respectively). In contrast, only 34% of TRs with motifs 10 bp or larger are isolated TRs, while nearly half (46%) are located within complex VCs (**Figure 5A**). We also observed a higher proportion of isolated TRs in coding or untranslated regions (94%) relative to intronic, promoter, or intergenic regions (88%). Conversely, a higher percentage of TRs in intronic, promoter, or intergenic regions lie within complex VCs (5%) compared to those in coding or untranslated regions (1%) (**Figure 5B**). Finally, we found that TRs within VCs tend to have lower mappability (as defined in **Table S3**) while TRs with high mappability are nearly all isolated TRs (**Figure 5C**).

**Figure 5.**
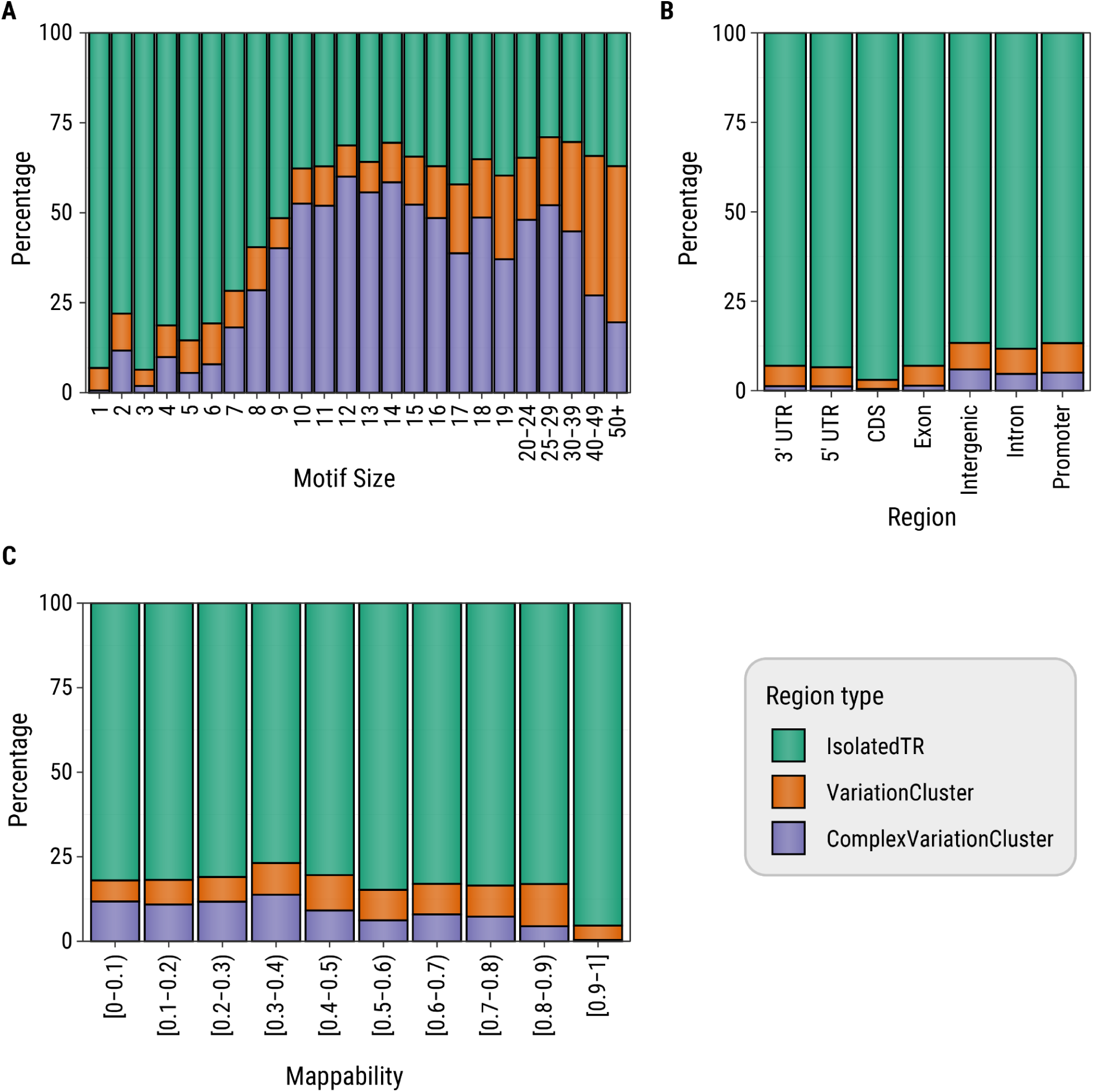
Percentage of tandem repeat (TR) loci in each variation cluster (VC) status group as a function of (A) motif size (in bp), (B) Gencode genomic region, and (C) mappability of tandem repeat locus and flanks. Isolated TRs are TRs that do not lie within a VC, complex VCs are VCs that contain five or more TRs.

We also assessed whether TRs or VCs occur more frequently near other repetitive elements, such as long or short interspersed nuclear elements (LINEs or SINEs), by comparing the proportion which overlap or lie within 100 bp of the eight most frequent major classes of repetitive elements in the UCSC Genome Browser RepeatMasker annotation track^47^ (**Figure S6A**). We also generated a set of 1,000,000 randomly selected genomic regions with lengths sampled from the empirical distribution of TR lengths to evaluate the background rate of overlaps. As expected, TRs overlap simple repeat elements as well as SINEs^48,49^ at a substantially higher frequency than random regions. SINE elements overlap isolated TRs and small VCs around twice as often as complex VCs, which are similar to the background rate (∼0.2). In contrast, isolated TRs and VCs overlapped LINE elements at a rate similar to the background, whereas complex VCs were low. TRs, VCs and complex VCs exhibited higher overlap with low complexity regions than background, with the strongest enrichment for complex VCs, with over twice the proportion of overlaps (∼0.1) than the next highest group. As TRs tend to cluster together, we examined whether the pattern of overlaps between TRs and repetitive elements varied depending on the number of nearby TRs (defined as the number of TRs separated by no more than 6 bp of sequence). As the number of nearby TRs in the region increased, the proportion of overlaps with simple repeat elements and low complexity regions increased, while the overlap with LINEs and SINEs tended to decrease (**Figure S6B**).

Additionally, we performed VC enrichment analysis by picking random samples containing from 1 to 90 genomes from our dataset. Then we calculated VCs for each set of samples and compared each pair of VCs by calculating the Jaccard Index (total span of the intervals belonging to both sets of VCs divided by the span of their union) between them. This analysis showed that the set of VCs becomes quite similar at the 10 sample mark (Jaccard Index > 0.8; **Figure S7**), suggesting that many VCs are common (note however that rare and population specific variation may occur within common VCs).

### Efficient Motif Sets

To further characterize sequence composition of TR loci in our catalog, we generated efficient motif sets for our catalog of 4,863,041 TRs at a compression level of q=0.1, following the vamos protocol.^24,50^ After filtering out loci that overlapped centromeres, had more than 500 unique motifs, or where the locus itself or its surrounding variation cluster spanned more than 10 kb, 4,734,075 loci remained. Among these, 2,138,077 loci (45%) had more than one unique motif but only 280,636 (6%) had more than one efficient motif. As shown in **Figure S8**, the replaced motifs were rare by counts among assemblies, indicating that efficient motifs effectively capture the primary sequence compositions in TRs.

### Using the TRExplorer Catalog

TR genotyping analyses involving catalog-based tools for short-read and long-read data can be divided into three steps: 1) catalog selection 2) genotyping, and 3) downstream analysis. In this section, we provide practical recommendations for these steps when using the TRExplorer catalog and companion resources. For the first step–catalog selection–the TRExplorer catalog provides a starting point or superset from which researchers can select the subset of TR loci most relevant to their study goals. While it is possible to genotype all 4.86M loci, this is unlikely to be useful or cost-effective for most studies. Instead, we recommend using the multi-faceted filtering and search functionality provided by the TRExplorer portal (**Figure S9**) to first select locus subsets based on the comprehensive annotations available, and then, using the “Export to” feature, download the selected loci in the format required by the TR genotyping tool of choice. Examples of filtering criteria that are likely to be generally applicable include selecting TR loci that A) exclude some motif sizes such as homopolymers, dinucleotides, and/or large VNTR motifs B) have motifs like those at known disease-associated loci; CAG, CCG, GAA, AAAAT, etc. C) are polymorphic in the HPRC or AoU phase 1 datasets, for example those with standard deviation (σ) ≥ 0.2, though the exact threshold should be study-specific D) do not reside within large variation clusters E) are not intergenic.

The TRExplorer v1 catalog contains 18,340 (0.4%) instances of overlapping locus definitions. We chose not to remove, deduplicate, or merge these definitions because different users may have different preferences on how to handle such loci. Therefore, in the “Export to” dialog, the TRExplorer portal provides the option to include overlapping definitions as separate loci, merge them, or export only one of the loci, prioritized by locus length or purity.

For the second step–TR genotyping–tool accuracy and compute costs are key considerations. While assessing tool accuracy is beyond the scope of this manuscript, we have evaluated genotyping costs for the full catalog as well as several useful subsets (**Table S4**). The full catalog contained all 4,863,041 loci. Subset 1 reduced this to 1,662,481 loci by excluding all homopolymer and dinucleotide motifs, along with all intergenic loci. Subset 2 further filtered subset 1 to the 240,163 loci that were polymorphic in the HPRC256 dataset, defined as having a population allele frequency distribution standard deviation ≥ 0.2. Subset 3 further filtered subset 2 to the 33,546 loci that had one of the following 14 normalized motifs seen at known disease-associated loci: CAG, GCC, GAA, GTC, CAGG, TTTG, GGGCC, ATTCT, GGCCCC, GGCCTG, GGCGCGGAGC, CGCGGGGCGGGG, CCTCGCTGTGCCGCTGCCGA, CCTCATGGTGGTGGCTGGGGGCAG. We selected a 30x-coverage short-read Illumina genome and a 30x PacBio HiFi genome from HG002 as our test samples. Then, we ran two representative short-read tools–HipSTR v0.7^1^ and a modified version of ExpansionHunter v5 (**Code Availability**),^31^ as well as two long-read tools–TRGT v5.0.0^18^ and LongTR v1.2^17^ on the catalog and each subset. We ran these analyses on standard Google Compute Engine virtual machines via the Hail Batch job running platform.^51^ When running on non-preemptible virtual machines that cost ∼$0.05 per CPU per hour, we found that genotyping subset 3 (34k loci) cost a total of between $0.02 and $0.10 per sample, subset 2 (240k loci) cost between $0.06 and $0.20, subset 1 (1.7M loci) cost between $0.18 and $0.41, while the full catalog (4.86M loci) could cost as much as several dollars per sample (**Table S4**). These costs are expected to vary across cloud platforms and with a variety factors like disk I/O speeds, whether preemptible machines were used, and the specific loci being genotyped, since loci in some genomic regions accumulate excessive read depth and are therefore much slower to genotype than the average locus. However, this analysis provides representative ballpark estimates of current costs.

For the third step–analysis of results–the TRExplorer portal provides a growing number of annotations for interactive exploration or download. **Table S3** lists the annotations provided with the v1.0 catalog release. As an illustration of their utility, we used the annotations to summarize basic properties of known disease-associated loci (**Table S2**) and to reproduce Figure 2 in Danzi et al. 2025^52^ which shows that most of the known disease-associated loci are some of the most polymorphic in the genome (**Figure S10**). We point users to the TRExplorer portal for an up-to-date list of annotations, including the latest population allele frequencies, and annotations derived from recent publications such as the Tanudisastro et al. analysis of sc-eTRs^53^ and the Manigbas et al. 2024 TR PheWAS in UK Biobank.^11^ All annotations are available via the Downloads page or, equivalently, via the “Export to” feature above the search results table (**Figure S9**).

## Discussion

Advances in sequencing technologies and genotyping tools may soon make TR analysis a standard part of rare and common disease discovery pipelines alongside SNVs, indels and SVs. We expect that this will involve a transition from the current variety of highly custom, specialized approaches to a small number of well-established methods and repeat catalogs. The lack of consensus around TR catalog design in particular may become increasingly problematic as population databases and other large-scale surveys of TR variation are made available. Our study is therefore an effort to build consensus by bringing issues and trade-offs involved in TR catalog design to the foreground.

While existing genome-wide TR catalogs^16,23,30,34^ have enabled a range of important discoveries, they are still primarily designed for specific tools and sequencing technologies that make them suboptimal choices for population resources of TR variation. Although newer catalogs^4,29,39^ cover a large fraction of repeats in the human genome, they use permissive locus definitions that include significant amounts of non-repetitive sequence. Such catalogs cannot be easily applied to studies based on short-read data which will remain important due to the wide-spread adoption and larger sample sizes of these datasets. Many older catalogs developed for short-read tools do support repeat copy number analyses in both short-read and long-read data. However, these catalogs remain incomplete, missing a substantial fraction (30% or more) of the polymorphic STR loci in our test sets.

We therefore developed a catalog that incorporates TRs identified in the reference genome as well as polymorphic TRs detected through cohort-based analyses, ensuring comprehensive representation of a wider range of TR loci that are amenable to both short-read and long-read analysis. To simplify adoption, we provide the catalog in the different formats expected by short-read and long-read tools. Additionally, we annotate the TR loci with their gene regions, population allele frequencies, and other properties, and then make this data available for interactive exploration, visualization, filtering, and export through an online portal at trexplorer.broadinstitute.org.

Our catalog stratifies TRs into two groups: isolated TRs and variation clusters. This second group is derived using a novel tool that extends the boundaries of TRs located within broader regions of variation. This is similar to how HipSTR and LongTR dynamically extend the repeat boundaries when variation is detected close to the original repeat region. While the HipSTR/LongTR approach improves genotyping accuracy, it can lead to discordant repeat definitions when these tools are applied to different sample batches because the boundaries are adjusted at genotyping time. In contrast, defining variation cluster boundaries a priori with our new tool vclust facilitates comparisons across studies/batches in downstream analyses.

We hope that the TR catalog and the computational tools introduced in this paper will serve as a starting point for a dynamic community resource that will address the needs of most research groups involved in profiling variation in TR regions. The adoption of a shared catalog across TR studies and population databases will help avoid future challenges in the reuse and interpretation of TR analysis results, including population allele frequencies and pathogenic thresholds.

Going forward, we will extend the catalog by adding TR loci not captured by our current approach. This includes a broader set of polymorphic VNTRs and interrupted repeats. For each locus, our catalog currently specifies only the TR motifs present in the reference genome, it will be important to supplement our TR definitions with motif sets observed in other genomes.^24,50^ Future work will characterize the accuracy of TR genotypes across loci, tools, and sequencing technologies to identify loci that can be accurately genotyped using short-read tools as well as those loci that can only be resolved using long reads. The catalog would also benefit from annotations related to methylation states, interruption patterns, somatic mosaicism, and most importantly, allele frequencies derived from larger and more diverse populations, along with genotype quality scores which indicate where allele frequencies may be inaccurate.

In conclusion, repeat catalogs are a critical part of TR analysis and poorly constructed catalogs can lead to low sensitivity by missing relevant loci as well as increase the likelihood of genotyping errors through the use of locus boundaries that are suboptimal for existing tools. In this paper we discussed issues and best practices related to catalog design. Also, we proposed a comprehensive catalog that can serve as the basis for short- and long-read analyses, thereby promoting consensus across studies and population resources.

## Methods

### Catalog comparison

All catalog comparison steps were implemented in the trexplorer-catalog GitHub repo compare_catalogs.py script. Prior to comparison, any compound locus definitions were split into their constituent TR loci, each of which had a single chromosome, start (0-based), end, and motif. For example, the compound definition of the *HTT* locus described by the expression (CAG)*CAACAG(CCG)* would be split into two separate definitions with the motifs CAG and CCG. For the Vamos v1.2 catalog, the first motif was selected from each motif set. The PlatinumTRs v1.0 TRGT catalog contained compound locus definitions that typically spanned multiple adjacent TRs without specifying the boundaries between them. To use the same approach for all catalogs, we attempted to split these compound definitions into their constituent TRs by locating stretches of perfect repeats of each specified motif within the reference sequence, as implemented in the str-analysis repo convert_trgt_catalog_to_expansion_hunter_catalog.py script. For 296,782 out of 7,722,729 locus definitions (3.8%), the exact boundaries of constituent TRs couldn’t be resolved in this way due to interruptions in the reference sequence, so we excluded them from comparisons and statistics.

In Table 1, comparison of each catalog with the test set of polymorphic STRs was performed by converting the catalogs to BED format, then running bedtools v2.31.0^54^ using the following command:

`bedtools subtract -A -a {STR_test_set.bed} -b {catalog_i.bed} | wc -l`

We then divided the number of loci found to be unique to the STR test set by the overall size of the test set (250,262).

### Identifying polymorphic TR loci in 78 T2T assemblies

To identify polymorphic TRs within haplotype-resolved T2T assemblies from 78 individuals for use as our TR test set and as the 4th source of TR loci for our TR catalog (Table 2, row #4), we employed the algorithm described in Weisburd et al.^31^ Briefly, we performed assembly-to-hg38 alignments and variant detection for each individual by running dipcall v0.3 with default parameters.^55^ Then, we filtered high-confidence insertion and deletion variants to the subset that represented tandem repeat expansions or contractions. For an insertion or deletion allele to be considered a tandem repeat variant, its non-reference sequence plus its flanking reference sequences needed to consist of 3 or more repeats of some motif while spanning at least 9bp. We then merged per-sample TRs from the 78 individuals by running the merge_loci.py script provided in the str-analysis repo. The pipelines for running dipcall and generating the merged set of TR loci are implemented in the str-truth-set-v2 repo run_dipcall_pipeline.py and run_filter_vcf.py scripts.

### Detecting All Perfect Repeats in the Reference Genome

To identify uninterrupted TR stretches in the reference genome, we located all sequences that spanned at least 9bp and consisted of at least three repeats of any 1-1000 bp motif. We implemented this method in the colab-repeat-finder GitHub repo perfect_repeat_finder.py script, and ran it as follows:

`python3 perfect_repeat_finder.py --min-repeats 3 --min-span 9 --min-motif-size 1 --max-motif-size 1000 hg38.fa`

Initially, we tried using TRF v4.09.1 for this purpose by setting its mismatch and indel penalties to prohibitively high values (i.e. 107), and running it using the command:

`trf409.macosx chr22.fa 2 10000000 10000000 80 10 2 2000 -h -ngs`

However, when we compared TRF output for hg38 chr22 to that of perfect_repeat_finder.py, we found that TRF missed 1,355 (2%) out of the 67,638 perfect TRs detected by our approach. At the same time, our approach detected all perfect TRs identified by TRF. Loci missed by TRF had a variety of motifs and locus sizes, including the following examples: chr22:39,325,454-39,325,467 (GAG), chr22:22,778,396-22,778,417 (GATATA), chr22:50,530,014-50,530,023 (CGG), chr22:43,504,251-43,504,261 (TGA).

### Variation plots

A variation plot (as shown in **Figure 3**) is a visualization of a matrix whose columns correspond to bases in some genomic region and whose rows correspond to sequenced samples. The element (*i*, *j*) of the matrix is the fraction of alternative bases observed at position *j* in the sample *i*. The numerator of this fraction counts deleted bases, mismatched bases, and inserted bases (an insertion is assigned the reference position of its anchor base). The denominator is given by the number of HiFi reads that span the entire region. The maximum value of the fraction is set to 1.0. Note that the purpose of variation plots is to visually assess the presence of variation in a given region of the genome without fully resolving it. In particular, mismatches are not distinguishable from deletions and the length of the insertions cannot be ascertained. Plotting functionality within TRGT can be used for detailed visual analysis of variation in TR regions. Variation plots can be generated with the following script: https://github.com/PacificBiosciences/vclust/blob/main/utils/variation_plot.py

### Detection of variation clusters with vclust

To determine the variation cluster (VC) that a given TR belongs to, we implemented a method called vclust, available on GitHub (https://github.com/PacificBiosciences/vclust). The method calculates the probability that each base surrounding a given TR belongs to the VC. It then iteratively extends the boundaries of the VC upstream and downstream by one base pair at a time until the probability that each flanking base belongs to a variation cluster drops below 0.5 (**Figure S11**). To calculate the VC probability for a given base (**Figure S11A**), we consider variation in a window starting at that base and extending 150 bp away from the repeat (**Figure S11B**). We discretize variation for each base of the window into six categories according to the fraction of observed alternative bases (≤0.10, ≤0.25, ≤0.75, ≤1.50, ≤5.00, and >5.00) producing a 150 element *variation vector* with entries in {0, …, 5}. We then calculate the probability that the first base of the variation vector window *W* belongs to the variation cluster using *P*(*VC* | *W*) = *P*(*W* | *VC*)*P*(*VC*) / (*P*(*W* | *VC*)*P*(*VC*) + *P*(*W* | *REF*)*P*(*REF*)).

Here, the terms *P*(*W* | *VC*) and *P*(*W* | *REF*) represent probabilities that the first base of the window *W* belongs to a variation cluster or is outside of the variation cluster respectively. They are calculated according to **Figure S11D** and **S11E**. The distributions in **Figure S11D** and **S11E** are defined through the following analysis: We randomly picked 50,000 TRs from our catalog, and then extracted the upstream and downstream flanking sequences of these TRs from 100 HPRC^56^ HiFi samples. The resulting sequences were aligned to the corresponding segments of the hg38 reference genome. If the base adjacent to the TR (rightmost base for the upstream flank and leftmost base for the downstream flank) was soft-clipped in more than half of the samples, we designated it as being in a variation cluster. We used the corresponding (discretized) variation windows to create the distributions depicted in **Figure S11D, E**. The terms *P*(*REF*) and *P*(*VC*) are set to 0.58 and 0.42 respectively corresponding to the fraction of repeat flanks that were softclipped by at least one base in this analysis. A locus was annotated as a variation cluster if its boundary was extended by more than 5 bp in either direction. Overlapping variation clusters were then merged together.

### TRGT longest pure segment genotypes

Longest pure segments (LPS)^52^ within repeat alleles were calculated using the TRGT-LPS tool (https://github.com/PacificBiosciences/trgt-lps). Briefly, TRGT-LPS finds copies of a given motif using a hidden Markov model (see TRGT paper)^18^ and then calculates the longest stretch of consecutive copies of that motif, which is allowed to have at most one imperfect motif per 10 perfect motif occurrences.

### Assembly consistency analysis

The assembly consistency analysis was performed as follows. We concatenated TR allele sequences produced by TRGT with 250 bp flanks extracted from the reference genome. We aligned the resulting sequences to the HG002 genome assembly with minimap2 2.24-r1122.^57^ The edit distance between each allele sequence and the corresponding segment of the assembly was then used as a measure of consistency.

## Acknowledgments

This work was supported in part by the Victorian State Government Operational Infrastructure Support and Australian Government National Health and Medical Research Council Independent Research Institute Infrastructure Support Scheme. BW and HR were supported by the National Human Genome Research Institute (NHGRI) under grant U01HG011755. LH is supported by the National Cancer Institute (NCI) under grant F30CA284847. HZJ and MG were partially supported by the National Human Genome Research Institute (NHGRI) under grant 1R01HG010149. BG and MJPC are supported by R01HG011649.

## Data Availability

The Repeat Catalog is available on the Releases page of the trexplorer-catalog GitHub repo (https://github.com/broadinstitute/trexplorer-catalog).

The 100 HiFi samples from the Human Pangenome Reference Consortium are available under BioProject ID PRJNA850430 (https://www.ncbi.nlm.nih.gov/bioproject/730823) in SRA. TRGT allele calls in VCF format are available on the Downloads page of trexplorer.broadinstitute.org

## Code Availability

● https://github.com/broadinstitute/TRExplorer TRExplorer online portal client-side and server-side code
● https://github.com/PacificBiosciences/vclust variation cluster analysis tool
● https://github.com/PacificBiosciences/trgt-lps TRGT companion tool that processes TRGT allele sequences to detect the longest pure segments (LPS)^52^
● https://github.com/broadinstitute/variation-clusters documentation and tutorials for computing variation clusters
● https://github.com/broadinstitute/trexplorer-catalog scripts for generating the new repeat catalog described in this paper
● https://github.com/broadinstitute/colab-repeat-finder tool for detecting perfect repeats in nucleotide sequences
● https://github.com/bw2/ExpansionHunter This modified version of ExpansionHunter v5 introduces “optimized-streaming” mode which trades-off slightly lower accuracy in favor of increased speed and reduced memory usage when processing large (> 5,000 locus) catalogs.
● https://github.com/broadinstitute/str-analysis utilities for filtering, annotating, and converting TR catalogs to a variety of formats
● https://github.com/broadinstitute/str-truth-set-v2 pipelines for identifying TR variants through T2T-assembly-to-reference-genome alignment via dipcall, followed by filtering and post-processing steps^31^

## Supplementary Information

### Different definitions of the *PABPN1* locus

In the hg38 reference, the PABPN1 locus begins with six GCGs (chr14:23,321,473 - 23,321,490), followed by three GCAs (chr14:23,321,491 - 23,321,499), followed by GCG⋅GGG⋅GCT⋅GCG. This sequence can be concisely represented as GCG[6]GCA[3]GCG[1]GGG[1]GCT[1]GCG[1]. Brais et al. (1998)^58^ established that *PABPN1* expansions to 8 or more GCG repeats cause autosomal dominant oculopharyngeal muscular dystrophy (OPMD). Their study did not include the adjacent GCA[3]GCG[1] in the estimation of the pathogenic threshold, and thereby set locus boundaries at chr14:23,321,473 - 23,321,490. However, subsequent studies^9,59^ and resources such as GeneReviews and STRchive^60^ extended the boundaries to chr14:23,321,473 - 23,321,499, including GCA[3]GCG[1], and therefore adjusted the pathogenic threshold to 12 or more repeats. In the meantime, other resources such as the STRipy database^61^ and gnomAD continued to use Brais et al.’s original locus boundaries and pathogenic threshold of 8. While these different definitions describe the same biological reality, the inconsistency creates the potential for inadvertent errors in OPMD diagnosis.

## Figure Ss

**Figure S1:**
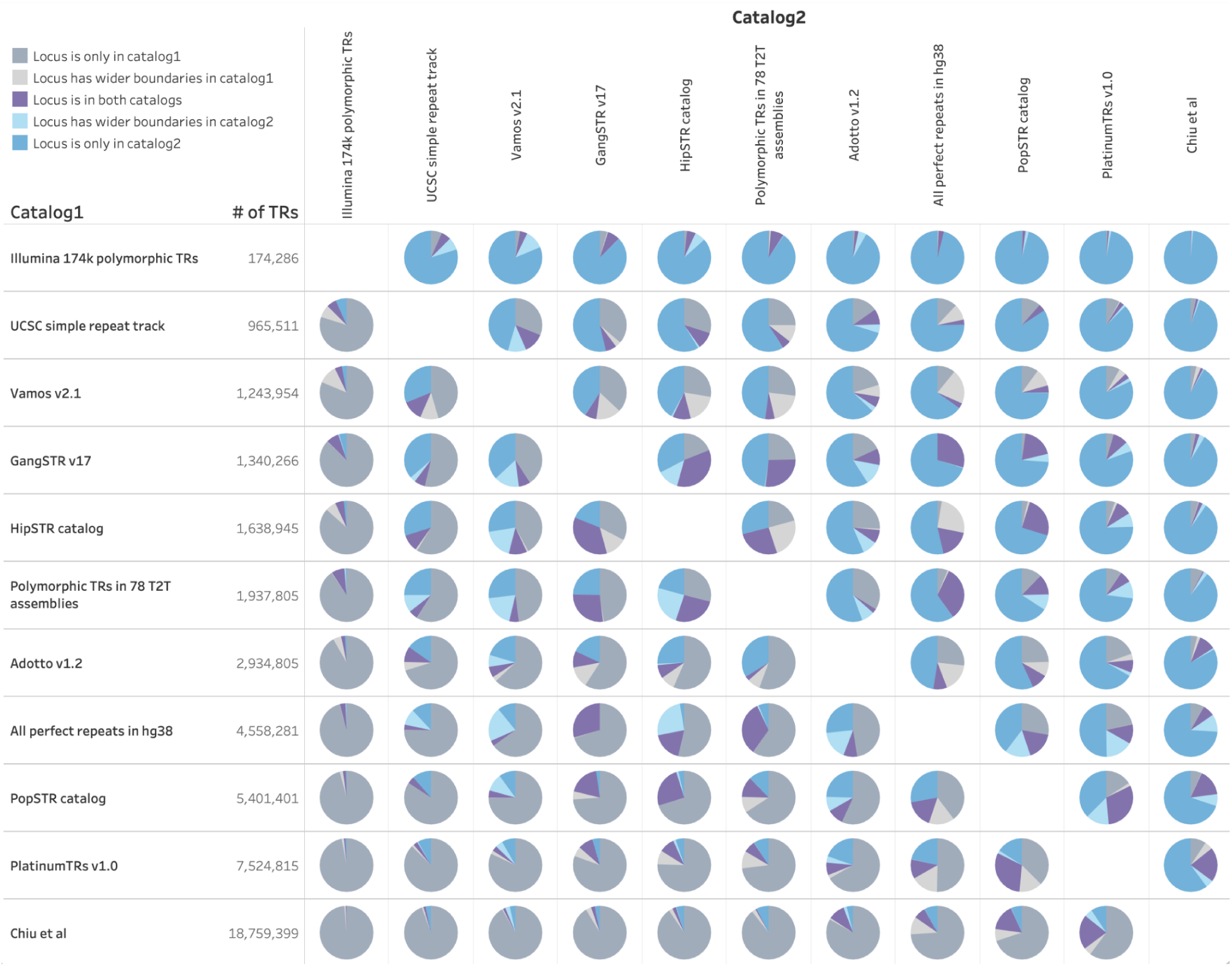
Pairwise comparison of TR definitions in 11 catalogs. A locus was considered to be in both catalogs (purple) if the two locus definitions overlapped each other by at least 5%, specified the same motif, and differed in size by less than one motif length. Motif cyclic shifts and reverse complements (i.e. CAG, AGC, GAC, CTG, TGC, GCT) were considered to be equivalent. Locus definitions with the same motif but wider boundaries in catalog 1 (light gray) or wider boundaries in catalog 2 (light blue) were counted separately.

**Figure S2:**
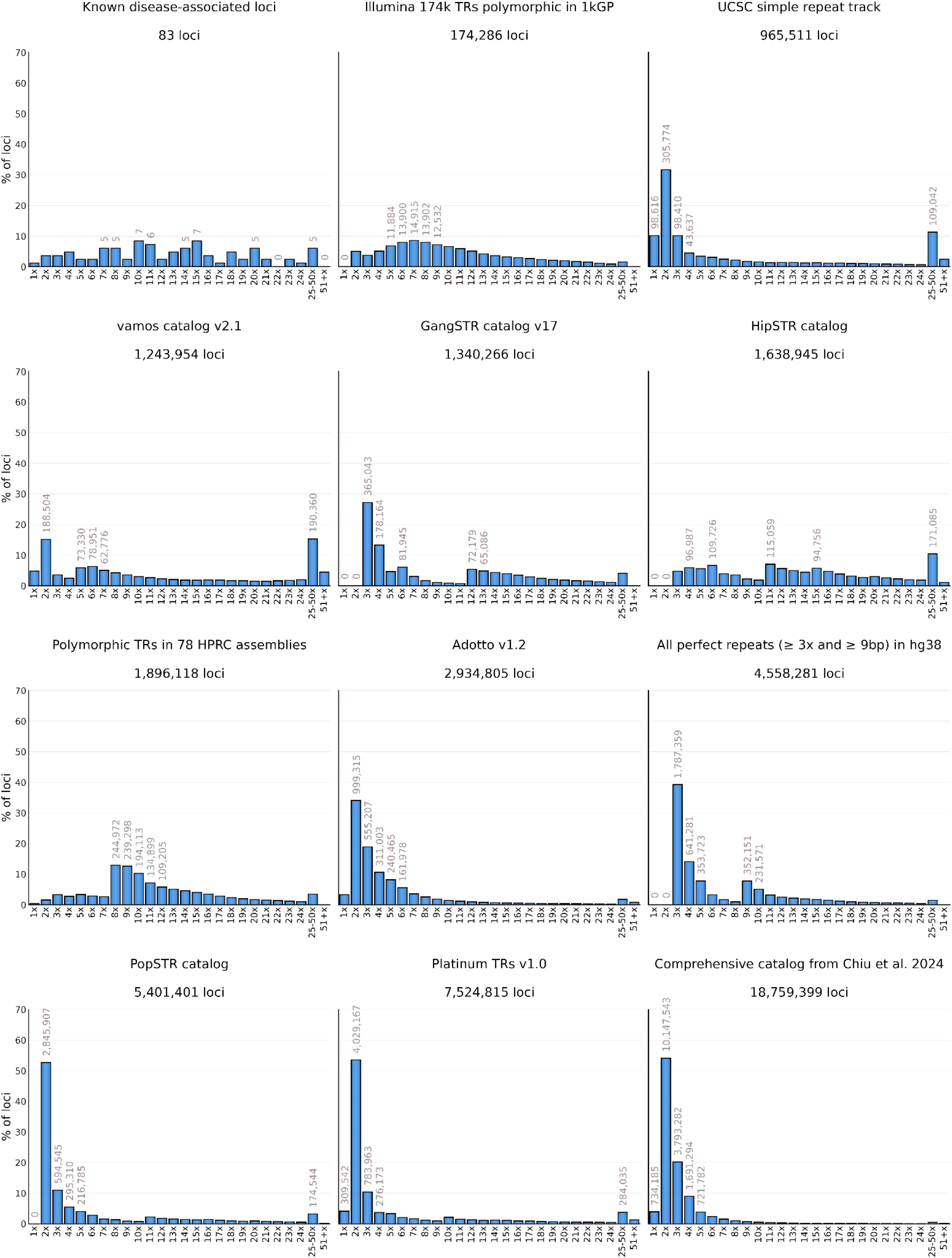
Reference locus size distributions in 12 TR catalogs. The horizontal axis represents the repeat copy number of each locus in the hg38 reference.

**Figure S3:**
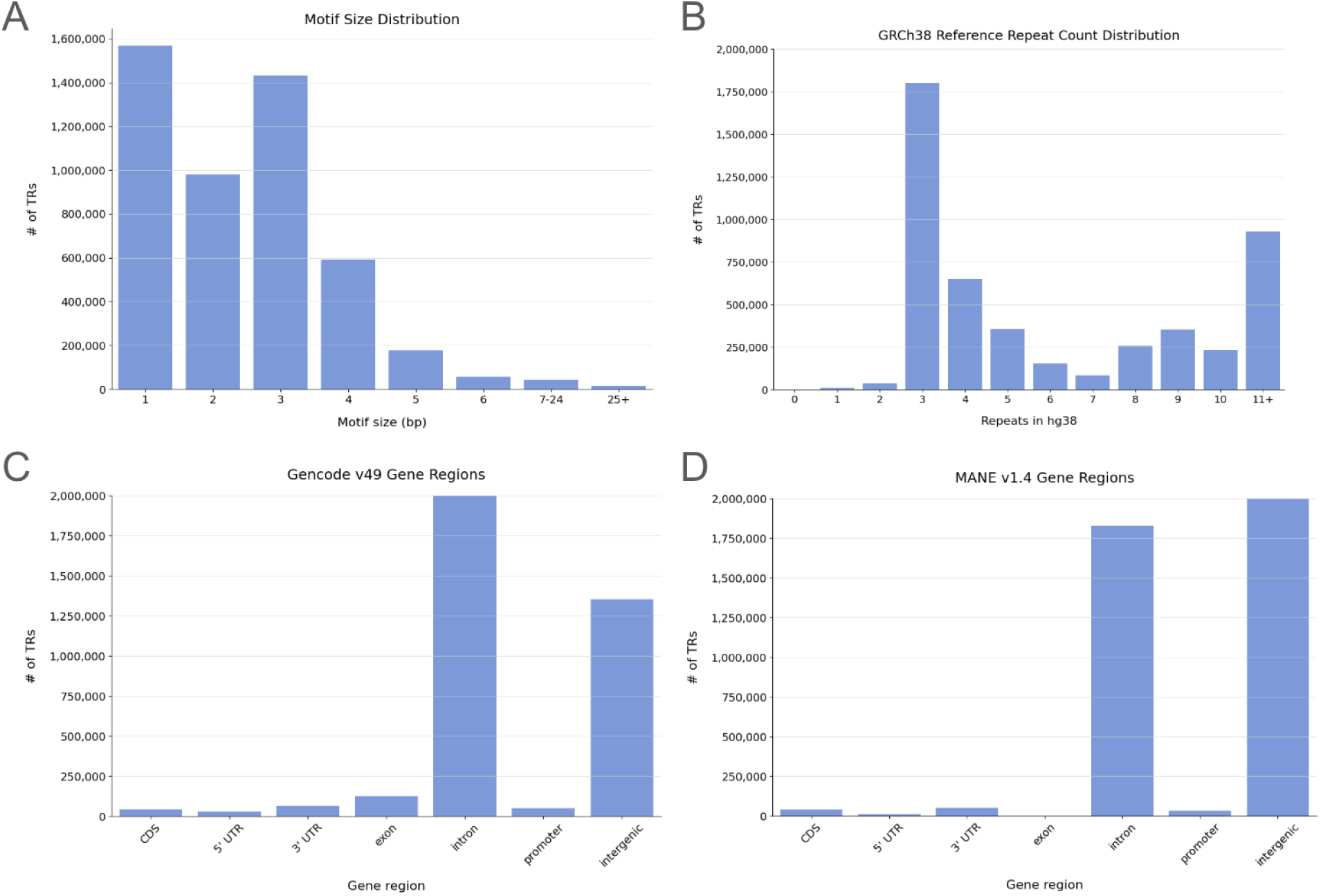
TRExplorer v1.0 catalog summary statistics. (A) Motif size distribution (B) Repeat copy number in the hg38 reference (C) Most significant gene region that overlaps the locus based on the Gencode v49 comprehensive annotation (D) Most significant gene region that overlaps the locus based on the MANE v1.4 annotation

**Figure S4:**
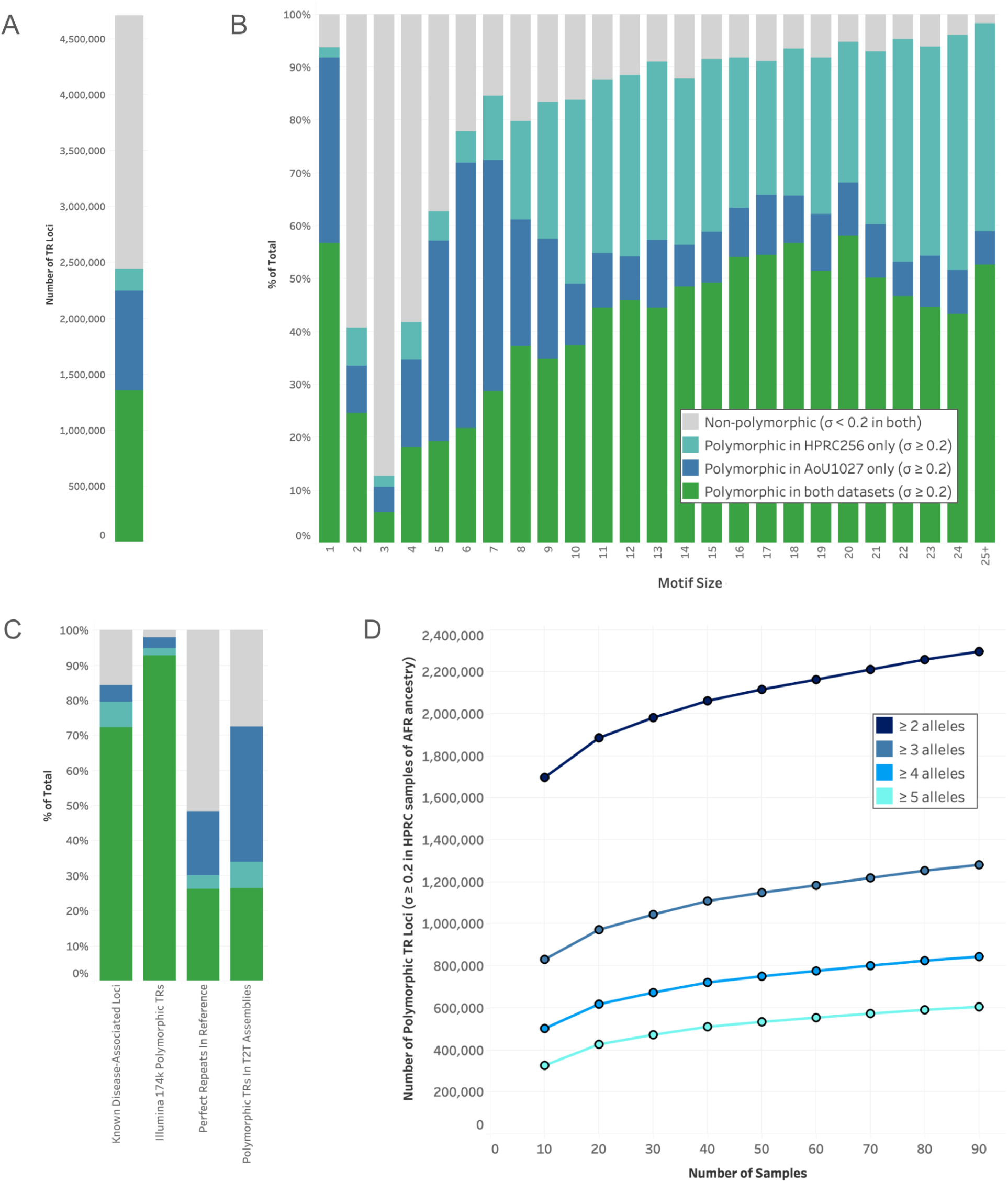
Polymorphism rates. Total counts (A) and fractions (B, C) of loci that are non-polymorphic (gray), defined as their allele size distributions having a standard deviation < 0.2 in both the AoU release 1 dataset of HiFi samples from 1,027 self-identified Black or African American individuals and in 256 HPRC HiFi samples from diverse ancestries. Fraction of loci that are polymorphic only in the HPRC samples (teal), fraction of loci that are polymorphic only in the AoU release 1 dataset (navy blue), fraction of loci that are polymorphic in both datasets (green) stratified by motif size (B) and source catalog (C). The number of loci that have no fewer than 2, 3, 4, or 5 observed alleles by repeat count in samples of African ancestry from the HPRC dataset, downsampled to 10, 20, .. up to 90 samples. TR repeat counts in both datasets were computed by running TRGT followed by TRGT-LPS to find the number of repeats in the longest pure segment in each allele.

**Figure S5:**
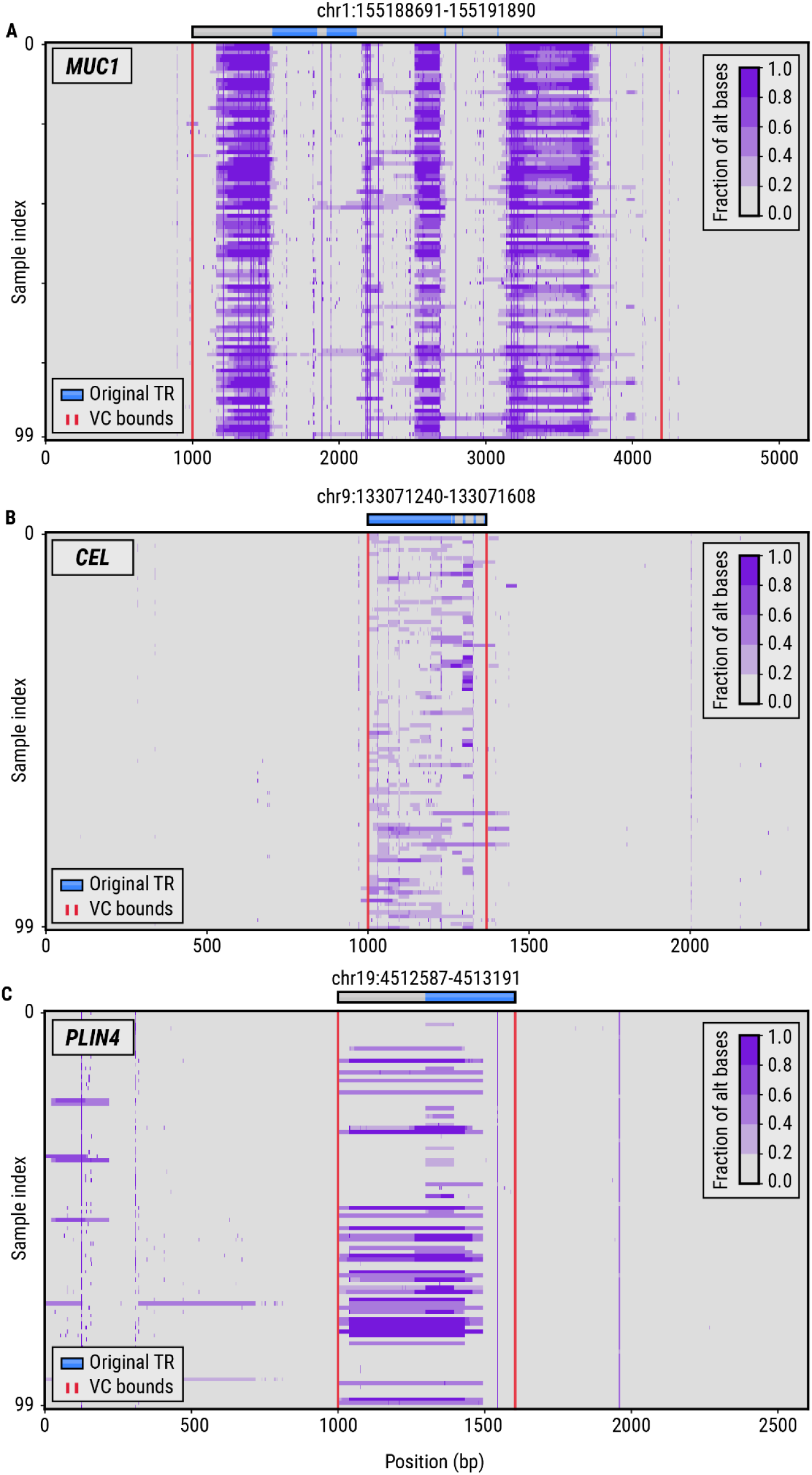
Variation plots showing the genetic variation in 100 HiFi HPRC samples around VNTRs in the (A) *MUC1*, (B) *CEL*, and (C) *PLIN4* genes. The blue bars denote the boundaries of the original repeat region while the gray bars depict the extension of the repeat region to a full-length variation cluster. The red vertical lines denote the boundaries of the variation cluster (VC) computed by the vclust tool.

**Figure S6:**
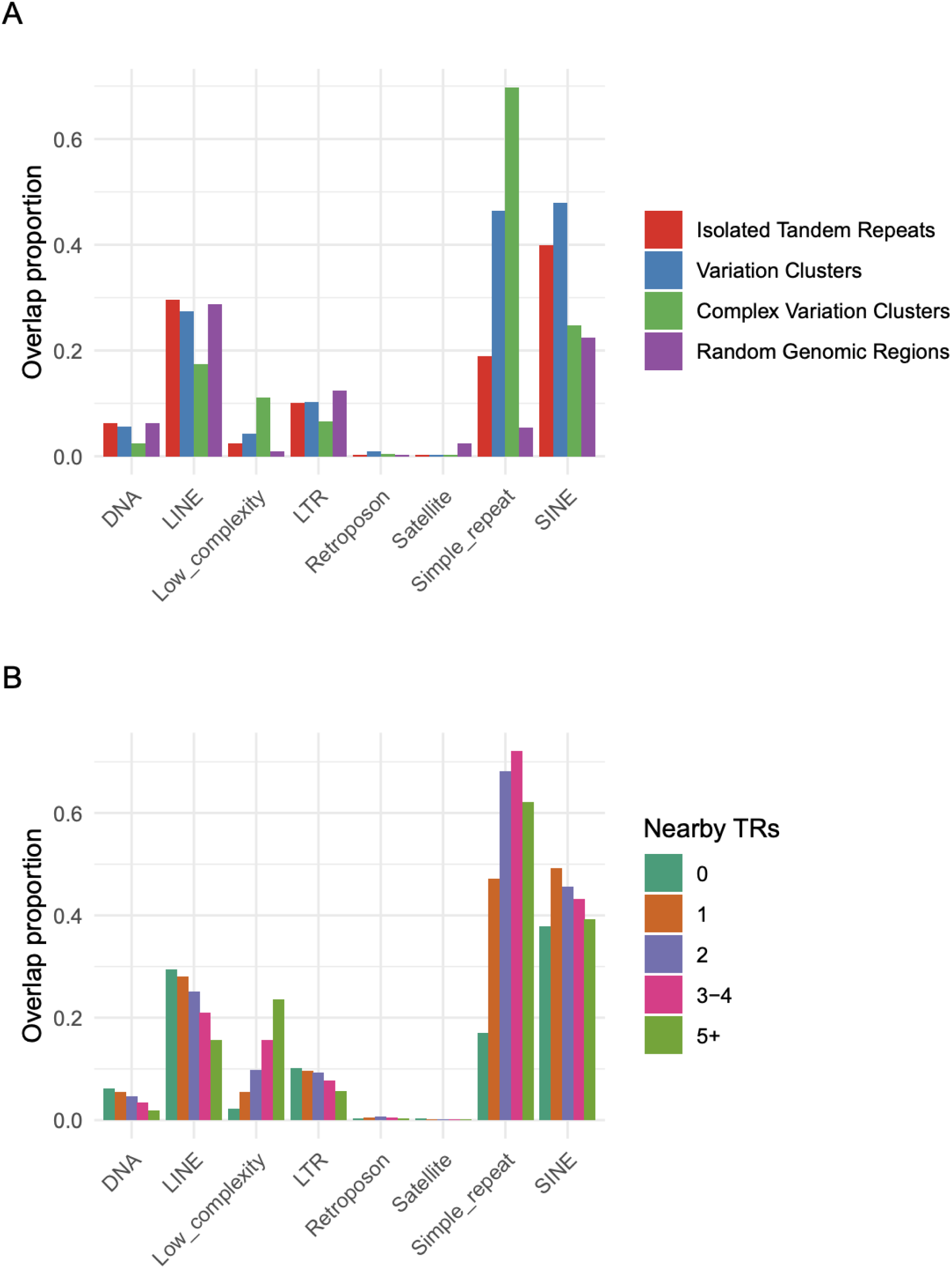
Overlap with repetitive elements. (A) Proportion of TR loci that overlap or lie within 100 bp of major types of repetitive elements in UCSC Genome Browser RepeatMasker track, compared to background rate for randomly selected genomic regions. TRs that overlapped multiple RepeatMasker classes were counted under each class. (B) Proportion of TRs that overlap repetitive elements, stratified by the number of nearby TRs (defined as TRs separated by no more than 6 bp of sequence). RepeatMasker abbreviations: DNA = DNA transposon, LINE = long interspersed nuclear element, LTR = long terminal repeat, SINE = short interspersed nuclear element.

**Figure S7:**
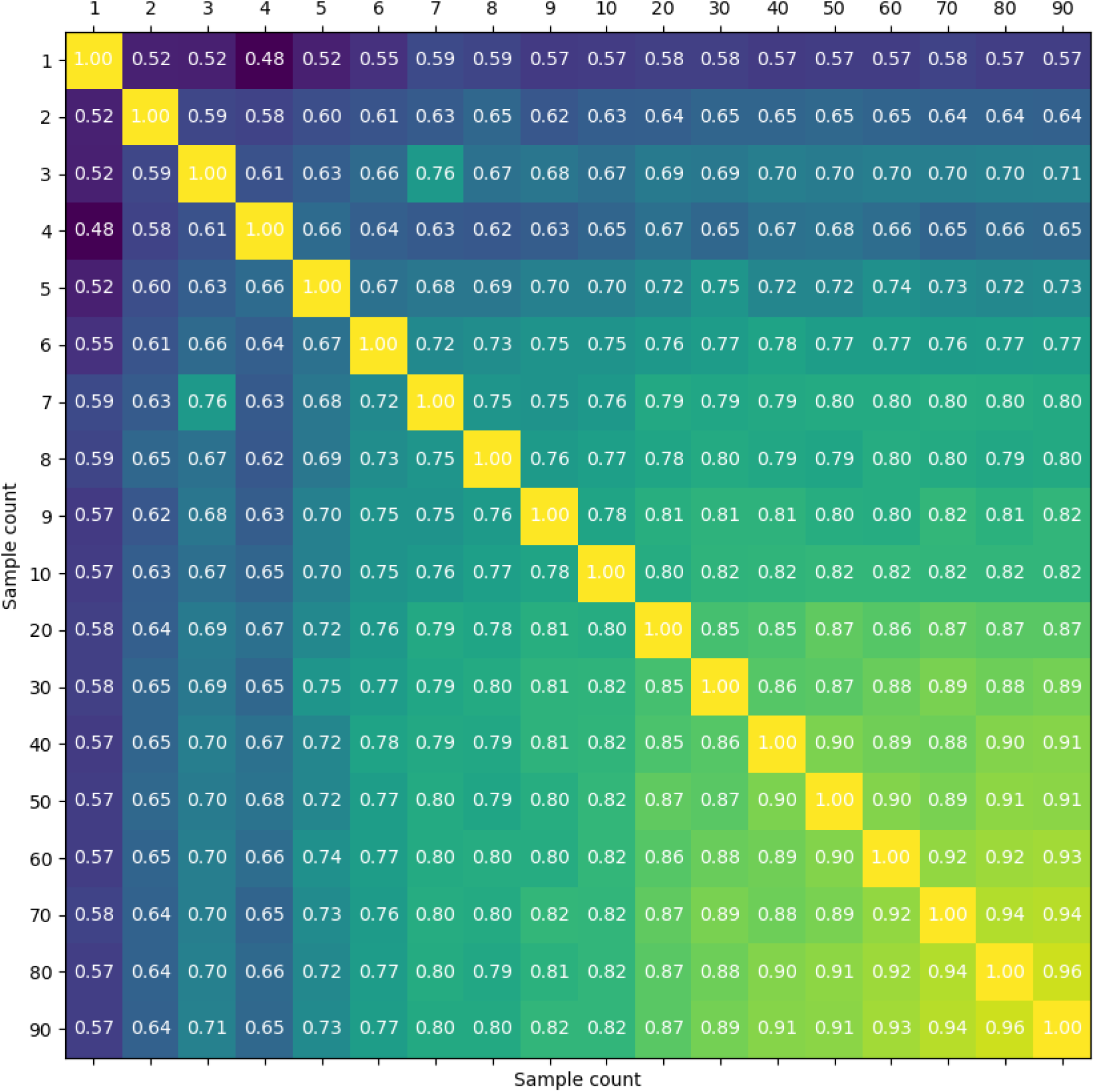
Variation cluster downsampling analysis. Jaccard Index from pairwise comparisons of VCs computed from different subsets of samples. For example, the value of 0.94 at x=70 and y=90 indicates that VCs calculated using a random subset of 70 samples had an average Jaccard Index of 0.94 when compared to VCs computed using a random subset of 90 samples.

**Figure S8:**
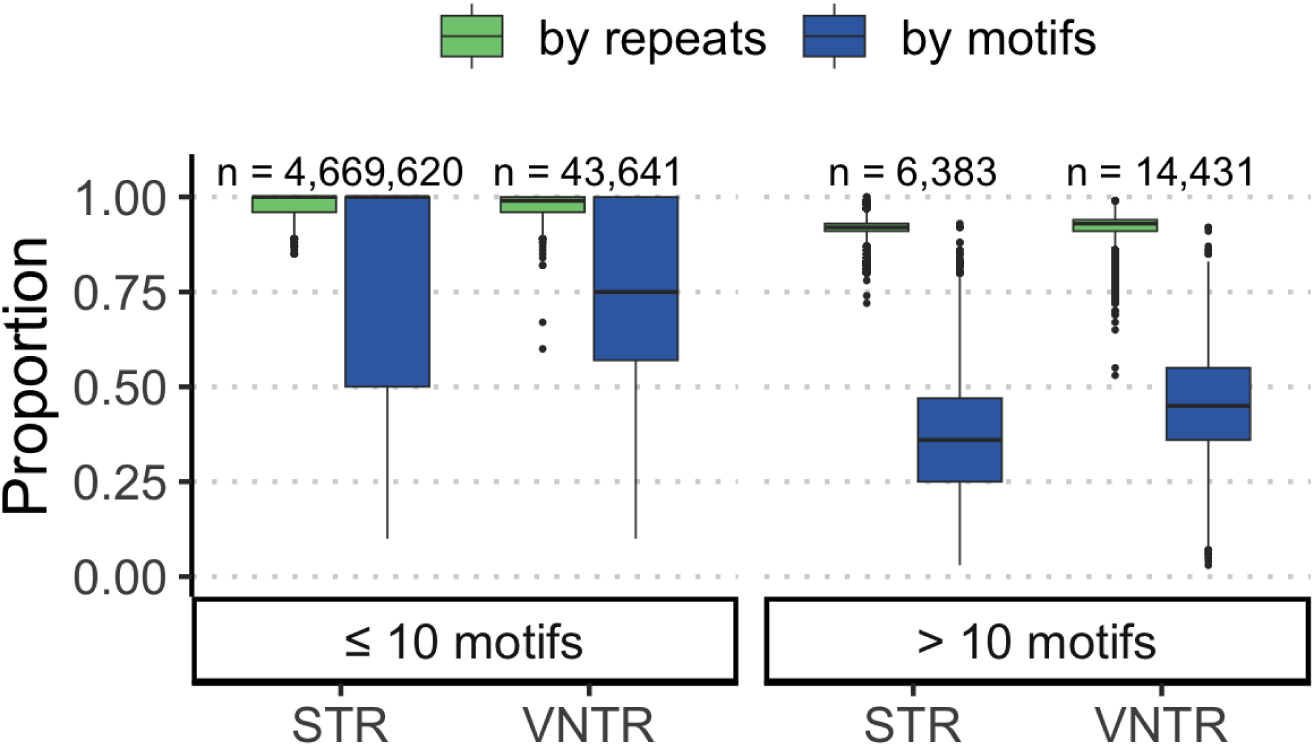
Proportion of efficient motifs. TRs were grouped by the number of unique motifs observed within the assembly repeat sequences at a locus (≤ 10 or > 10). Green boxes represent the fraction of assembly repeat sequences that are composed of efficient motifs (q = 0.1) while blue boxes represent the number of efficient motifs divided by the total number of unique motifs at a locus.

**Figure S9.**
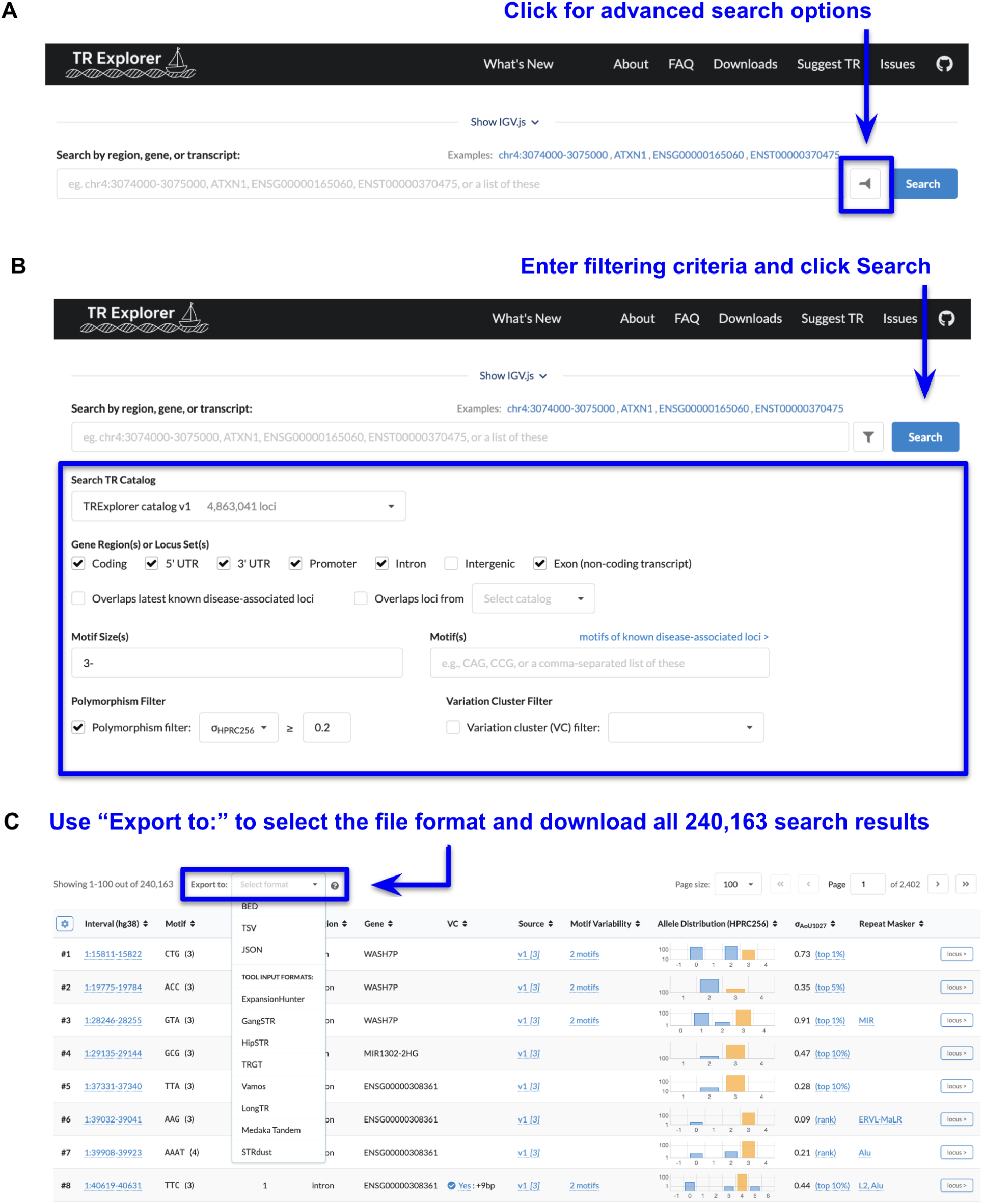
TRExplorer. online interface supports filtering and exporting a subset of the catalog by various criteria. This example demonstrates using (A) advanced search options to (B) specify the subset of TR loci that are not intergenic, have a motif size of 3bp or more, and are polymorphic–defined as having a population allele frequency distribution standard deviation ≥ 0.2 in the HPRC256 dataset. After clicking the Search button, the filtered catalog can be downloaded via the “Export to:” drop-down above the search results table (C).

**Figure S10:**
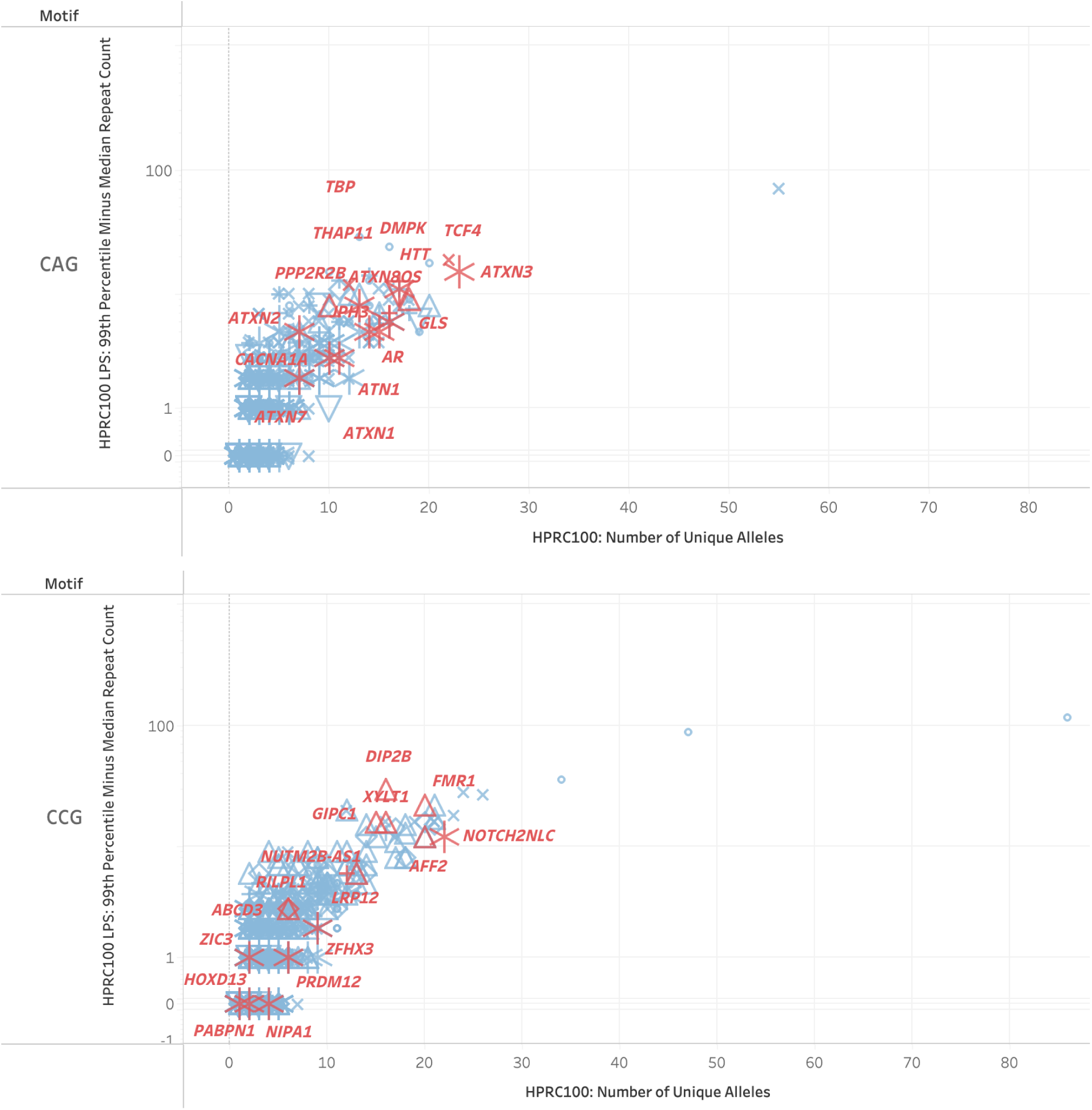
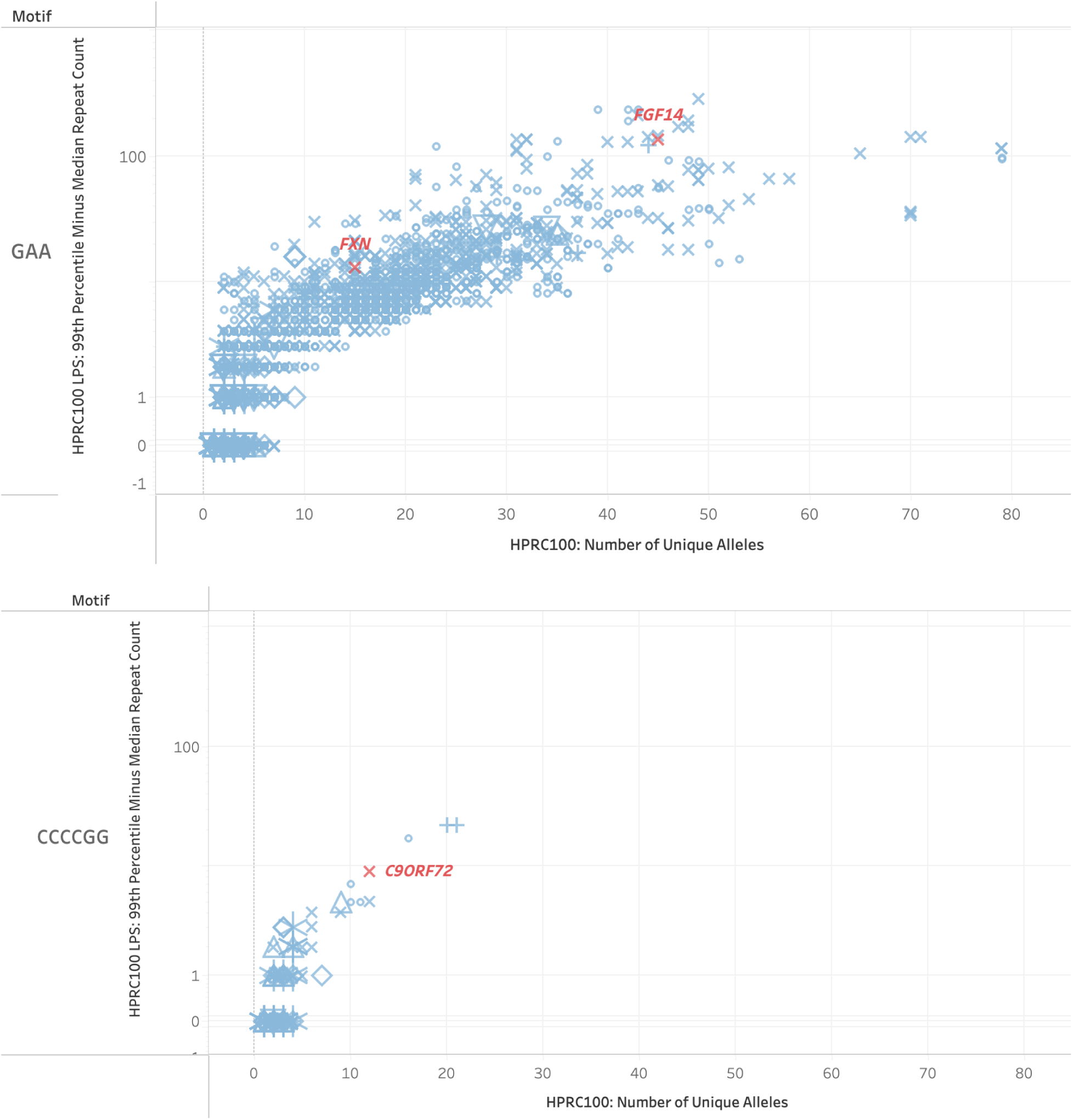
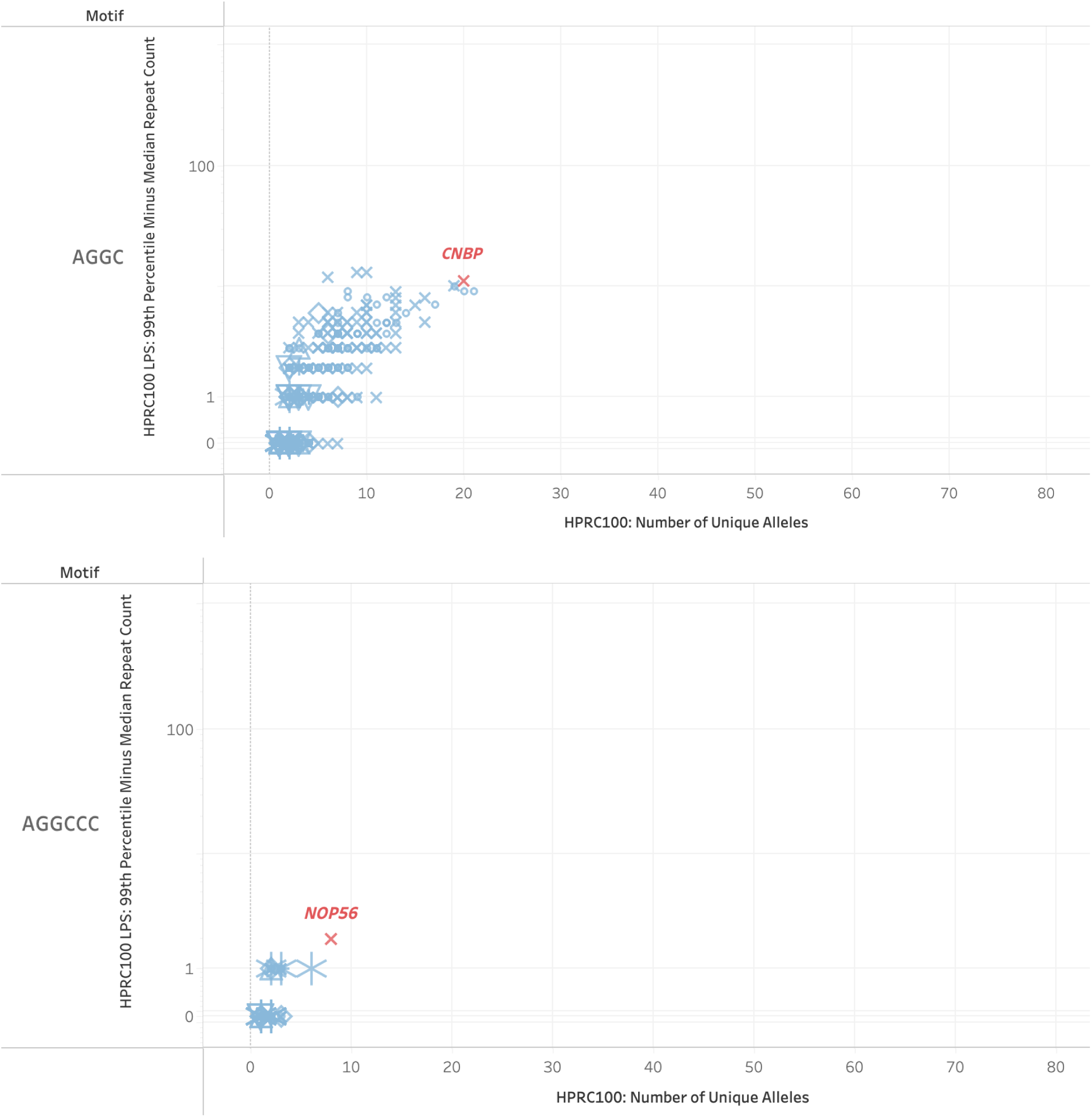
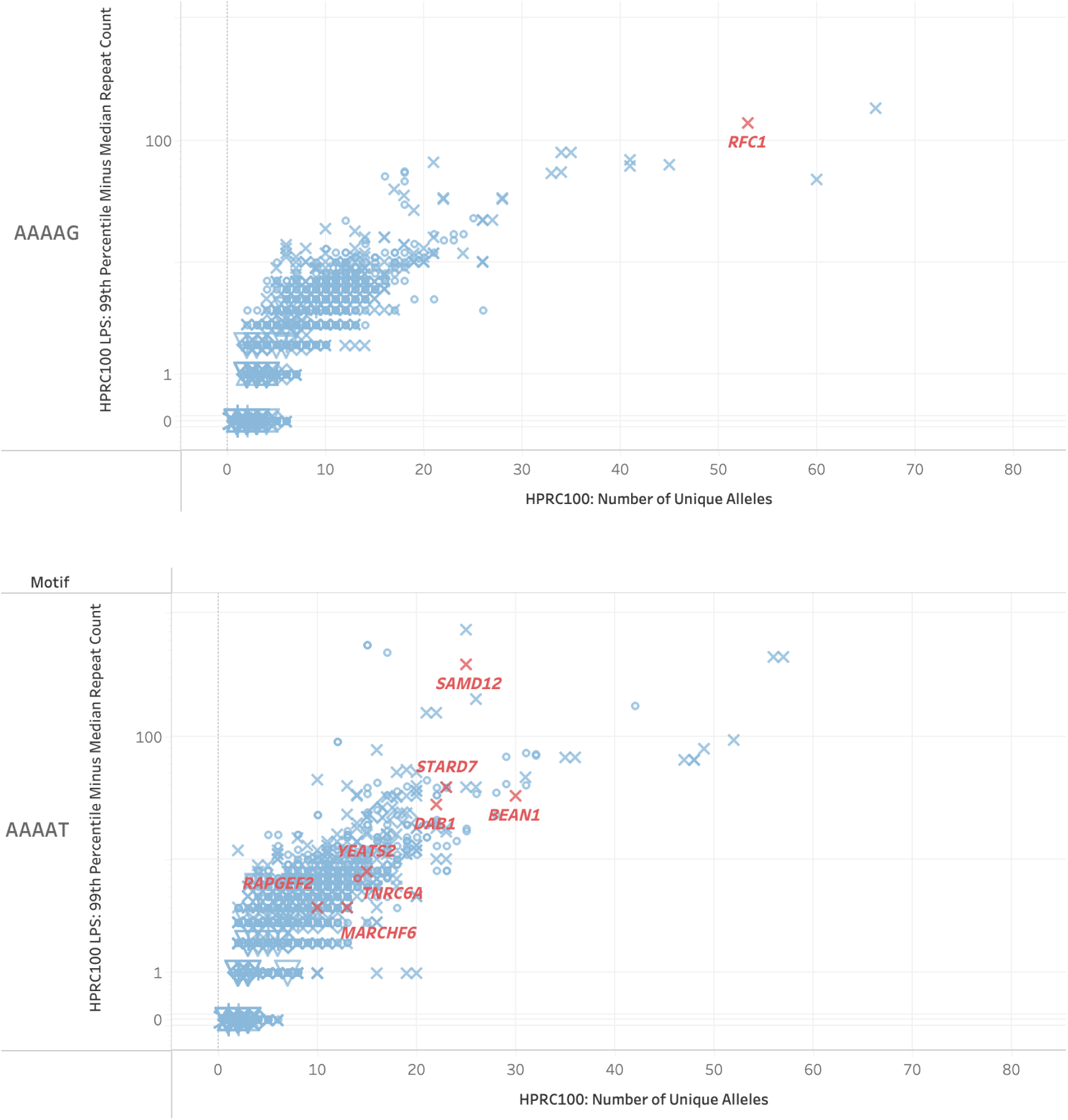
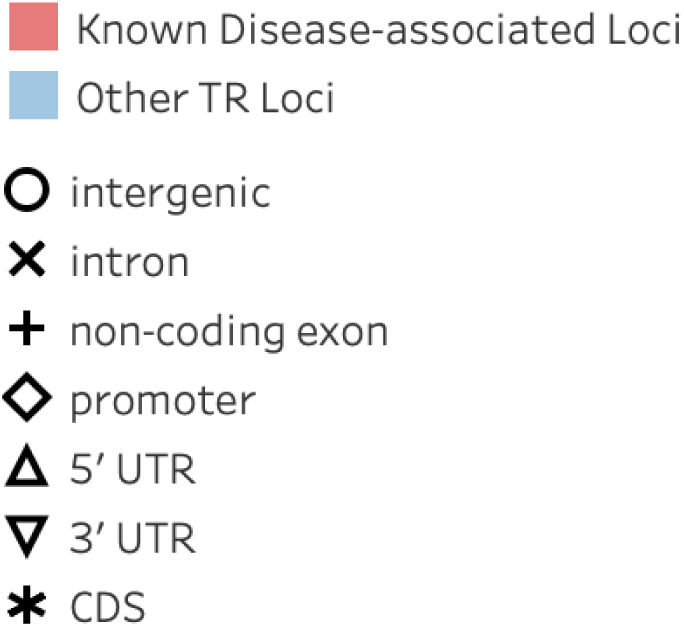
Polymorphism rates stratified by motif. Comparison of known disease-associated STR loci with all other loci in the catalog that share the same normalized motif (where shifted or reverse-complemented motifs such as CAG, AGC, GCA, TGC, etc. are considered to be the same). Each panel shows data for a different STR motif: (A) CAG (B) CCG (C) GAA (D) CCCCGG (E) AGGC (F) AGGCCC (G) AAAAG (H) AAAAT. Two metrics of polymorphism are shown on the x- and y- axis: the x-axis shows the number of observed TR allele sizes at a locus, while the y-axis shows the 99th percentile allele size minus the median allele size (in units of repeat counts) at a locus. The color is red for known disease-associated loci and blue for all other loci. Different shapes represent the gene region of each locus in Gencode v46. The polymorphism metrics were generated by running TRGT-LPS with the full TR catalog on 100 long-read HPRC samples from diverse populations.

**Figure S11:**
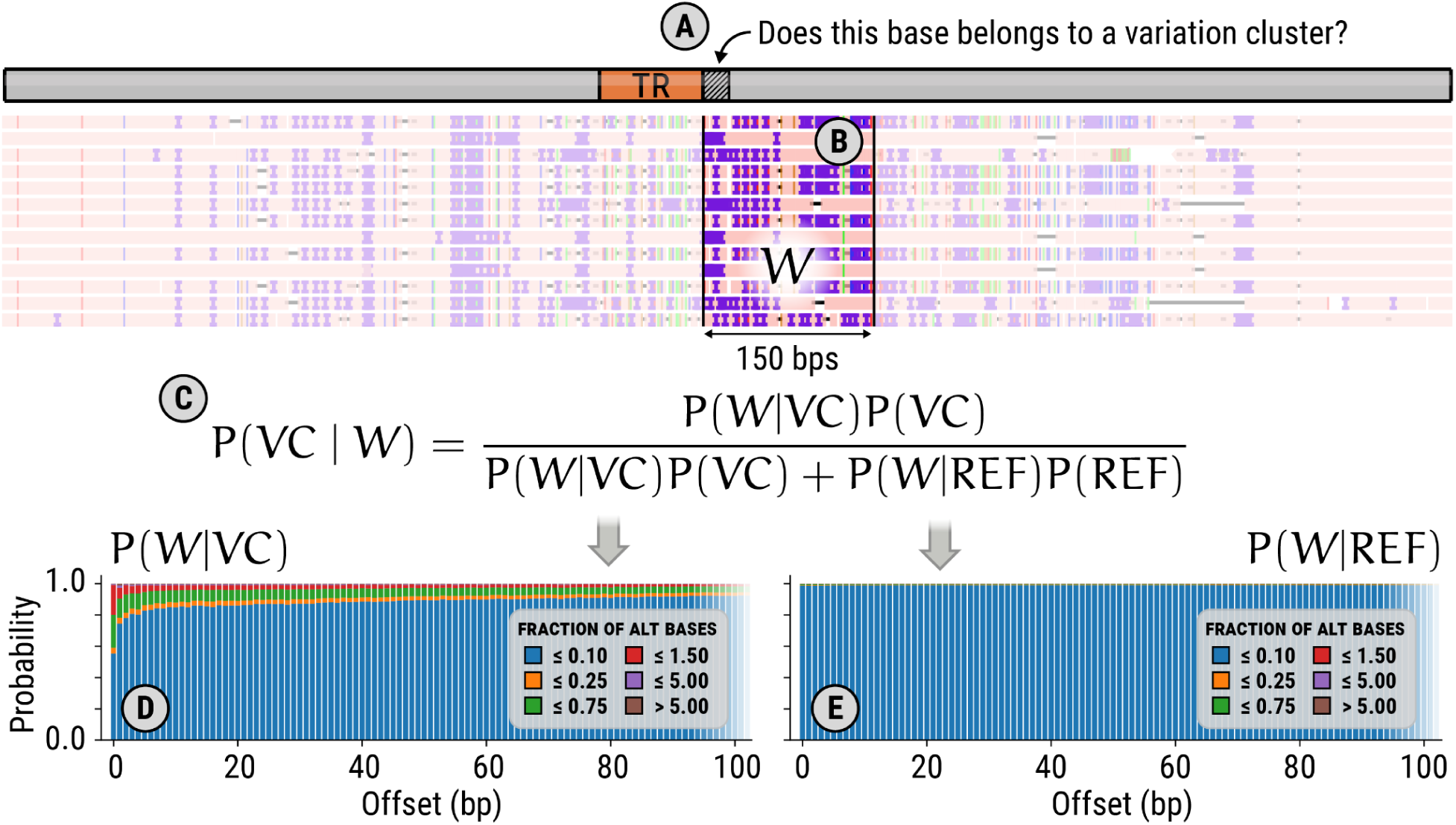
Computing variation cluster boundaries. (A) The base being evaluated is adjacent to the tandem repeat. (B) The 150bp variation window that starts at the base under evaluation and extends away from the repeat. (C) The equation used to evaluate the probability that the first base of the window *W* belongs to a variation cluster. (D) The model used to evaluate *P*(*W*|*VC*). (E) The model used to evaluate *P*(*W* | *REF*).

## Supplementary Tables

**Table S1: Summary of the methods used to generate existing TR catalogs.** This table shows the methods and parameters used to generate existing catalogs listed in Table 1.

**Table S2: Summary of annotations for known disease-associated loci.** This table lists the hg38 start (0-based) and end coordinates, motifs, gene regions based on Gencode v46, polymorphism rates (computed as the standard deviation of the TRGT-LPS repeat size distribution in 100 long-read samples from the HPRC), average mappability of the locus and flanking regions, whether the locus is embedded within a larger variation cluster (and if yes, how much larger it is than the TR locus), and finally, the number of other TR loci near this locus.

**Table S3: Per-locus annotations provided with the catalog.** Some annotations such as GencodeGeneName are only available for a subset of TRs. The six annotation categories are: Basic locus properties derived from the catalog itself or the reference genome (green), mappability (orange), variation clusters (blue), whether this is a known disease-associated locus (red), gene annotations (yellow), locus variability and population allele frequencies (purple).

**Table S4: Catalog genotyping costs.** Evaluation of TR genotyping runtimes and costs for the full catalog as well as subsets of different sizes in a 30x-coverage Illumina genome sample and a 30x PacBio HiFi genome sample from HG002.

